# Adenovirus E1A Binding to DCAF10 Targets Degradation of AAA+ ATPases Required for Quaternary Assembly of Multiprotein Machines and Innate Immunity

**DOI:** 10.1101/2020.12.16.423151

**Authors:** NR Zemke, WD Barshop, J Sha, E Hsu, JA Wohlschlegel, AJ Berk

**Affiliations:** Molecular Biology Institute, University of California, Los Angeles, CA 90095; Department of Cellular and Molecular Medicine, UCSD School of Medicine, La Jolla, CA 92093; Department of Biological Chemistry, David Geffen School of Medicine at UCLA, Los Angeles, CA 90095; Thermo Fisher Scientific, 355 River Oaks Parkway, San Jose, CA 95134; Department of Biochemistry and Molecular Medicine and the Norris Comprehensive Cancer Center, Keck School of Medicine, USC, Los Angeles, CA 90089; Department of Microbiology, Immunology, and Molecular Genetics, University of California, Los Angeles, CA 90095

**Author notes:** Corresponding author: A. J. Berk.

## Abstract

Adenovirus E1A early proteins modify host cell physiology to optimize virus replication. The N-terminal half of small e1a interacts with RB-family proteins to de-repress dNTP and DNA synthesis, and with p300/CBP to inhibit host anti-viral innate immune responses. These e1a N-terminal interactions activate a strong, late host anti-viral response due to stabilization and activation of interferon response factor 3 (IRF3). However, the C-terminal half of e1a inhibits this through interactions with three host proteins with seemingly unrelated functions. Proteomic analysis showed that all three C-terminal interactions are required for e1a-association into an ∼1 MDa multi-protein complex with scaffold subunits of a CRL4 E3 ubiquitin ligase and DCAF10, a presumed specificity subunit. This e1a-DCAF10-CRL4 prevents IRF3 stabilization indirectly by directing degradation of the essential AAA+ ATPases RUVBL1/2, subunits of several HSP90 co-chaperones required for quaternary assembly of cellular protein machines required for anti-viral defenses and responses to genotoxic and metabolic stress.

## INTRODUCTION

Human adenoviruses HAdV-2 and −5 are closely related DNA tumor viruses that infect epithelial cells lining the upper respiratory tract. To produce maximum progeny, adenovirus represses innate immune responses and forces these terminally differentiated, growth-arrested cells into S-phase to achieve high rates of viral DNA replication. E1A is the first adenovirus gene expressed following infection, generating two major protein isoforms during the early phase: large E1A (∼33 kD), which contains a transcriptional activation domain that activates transcription of viral early genes (*1*–*3*), and small E1A (“e1a,” ∼29 kD), which lacks the transcriptional activation domain but is sufficient to force G1-arrested cells into S-phase (*4*) and promote oncogenic transformation of cultured primary mammalian cells by cooperating with other oncogenes such as adenovirus E1B and G12V H-RAS (*5*–*7*). Conserved regions in e1a’s N-terminal half activate cell cycle genes by inhibiting RB-family proteins (RBs) (*8, 9*), and repress cellular differentiation and anti-viral defense genes by inhibiting the closely related major nuclear lysine-acetyl transferases p300 and CBP (*10*). While e1a’s C-terminal half also contains regions highly conserved between the ∼50 primate adenovirus species, it is unclear how these regions contribute to virus fitness.

Distinct peptides in the C-terminal half of e1a are bound by alternative cellular proteins (Figure S1). The paralogous FOXK1 and 2 transcription factors bind to a conserved serine/threonine rich region of e1a between amino acids 169 and 193 via their FHA (forkhead-associated) domains that bind phospho-Ser/Thr-containing peptides (*11*). DCAF7 associated with protein kinases DYRK1A, DYRK1B, and HIPK2 binds to amino acids 202 to 227 (*12*). And the paralogous transcriptional co-repressors CtBP1 and 2 bind to the conserved PXDLS CtBP-binding motif near e1a’s C-terminus (*13*). These three interactions are mediated through distinct e1a regions allowing for mutational disruption of any single interaction without interfering with the other two (*18*).

We reported previously on changes in host cell gene expression after infection of primary cells with HAdV-5 vectors expressing wt e1a or e1a mutants defective in binding either FOXK transcription factors (e1a mutant “FOXKb-”), DCAF7 complexes (e1a mutant “DCAF7b-“), or CtBP1 and 2 (e1a mutant “CtBPb-”). These vectors express normal levels of wt small e1a or these e1a mutants from the HAdV-2 E1A promoter/enhancer. The e1a C-terminal mutants contain two or four amino acid substitutions that inhibit detectable e1a binding to FOXK1/2, DCAF7 and its associated DYRK1A/B and HIPK2 protein kinases, or CtBP1/2 (*18*). Infection of G1-arrested primary human bronchial/tracheal epithelial cells (HBTECs) derived from the natural host tissue for HAdV-2 and −5, with vectors expressing each of the e1a C-terminal mutants activated expression of largely overlapping subsets of host cell genes, including ∼50 of the more than 200 genes classified as interferon-stimulated genes (ISGs) activated by all three of the e1a C-terminal mutants (*18*). Each of these e1a C-terminal mutants caused stabilization of IRF3, a sequence-specific DNA-binding activator of ISGs (*19, 20*), which reached high concentration, was transported into nuclei, associated with promoters and activated transcription of this subset of target ISGs (*18*).

Importantly, infection with the E1A deletion mutant *dl*312 (*21*) did not induce most of these ISGs, even though *dl*312 has normal virions (except for mutations in the viral DNA), and its viral DNA is transported into nuclei of infected cells using the normal HAdV-5 infectious process involving endocytosis, partial uncoating in early endosomes, transport of subvirion particles on microtubules to the microtubule organizing center near nuclear pore complexes, and finally transport of viral DNA coated with HAdV-5 protein VII through NPCs into the nucleoplasm (*22*). The observation that many ISGs were induced by the e1a C-terminal mutants, but not by infection with *dl*312, demonstrated the existence of an innate immune response triggered by mutant adenovirus e1a protein and not by detection of adenovirus nucleic acids exposed during the adenovirus infectious process. Note that during adenovirus infection, adenovirus DNA remains tightly associated with the highly basic internal virion protein VII until the viral genome is transcribed in the nucleus (*23, 24*). This may sterically block detection of Ad DNA by TLR9 in endosomes and by cyclic GMP-AMP synthase (cGAS) and other sensors for DNA in the cytoplasm (*25*). These observations led us to investigate how e1a C-terminal mutants induce IRF3 stabilization and how IRF3 stabilization is prevented by interactions between the C-terminal half of e1a and three cellular nuclear proteins that have no obvious related function.

Here we report that e1a inhibition of p300/CBP acetyl transferase activity induces IRF3 stabilization and activation. However, wt e1a counters this IRF3 stabilization by assembling a Cullin-based E3 ubiquitin ligase with specificity factor DCAF10, that stimulates proteasomal degradation of IRF3 indirectly. It does this by promoting degradation of RUVBL1 and RUVBL2, previously shown to be required for maximal activation of ISGs in response to interferon α (*26, 27*). RUVBL1 and 2 are essential subunits of several HSP90 co-chaperones required for the quaternary assembly of cellular multi-protein machines involved in anti-viral processes and responses to genotoxic and metabolic stresses. Our results reveal new mechanistic insights into how the activity of a viral protein can trigger an anti-viral defense response and how that same viral protein evolved to suppress this host cell defense in the ongoing “epic battle” between virus and host.

## RESULTS

### An antiviral innate immune response triggered by adenovirus e1a C-terminal mutants

To assess the influence of e1a’s C-terminal interactions on cellular gene expression we constructed HAdV-5 vectors that express wt e1a or an e1a C-terminal mutant with two or four amino acid substitutions that are defective for binding FOXK1/2 (“FOXKb-” = T229A, S231A, large E1A numbering), DCAF7 complexes (“DCAF7b-” = R258E, D271K, L272A, L273A), or CtBP1/2 (“CtBPb-” = PLDLS to ALAAA at amino acids 279-283) (Figure S1) (*18*). To inhibit expression of HAdV-5 genes other than wt e1a or these C-terminal mutant small e1as, these vectors did not express large E1A, the principal E1A transcriptional activator, because of a 9 bp deletion that includes the 5’ splice site required for producing the 13S E1A mRNA encoding the large E1A protein isoform. As a result, these vectors are defective for activating expression of the early viral mRNAs (*2*). RNA-seq from HBTECs 24h p.i. with these vectors identified 52 genes activated > 2-fold by all three e1a C-terminal mutants (FOXKb-, DCAF7b-, and CtBPb-) (*18*) (Figure 1). These genes are mostly reported interferon-stimulated genes (ISGs, > 90%) with established antiviral activities. Surprisingly, the majority of these genes (33/52) were not activated, or were activated much less (<2-fold), in response to viral DNA released or exposed during infection with the E1A deletion mutant *dl*312 (ΔE1A) (*21*) (Figure 1A, top cluster: e1a C-terminal mutant responsive genes). The remaining 19 of the 52 genes induced by infection with a C-terminal e1a mutant were also induced by infection with mutant *dl*312 which has a nearly complete deletion of E1A (*21*). Mutant *dl*312 is packaged into a normal HAd5 virion and introduces its packaged viral DNA into the host cell nucleus similarly to wt HAdV-5; however, *dl*312 does not express detectable viral proteins at 24h p.i. Therefore, these C-terminal mutant e1a proteins, and not viral DNA, triggers activation of these ISGs classified as “e1a mutant Responsive Genes” (Figure 1A). They include the well-studied ISGs *IFIT2, OASL*, and *MX1*.

**Figure 1.**
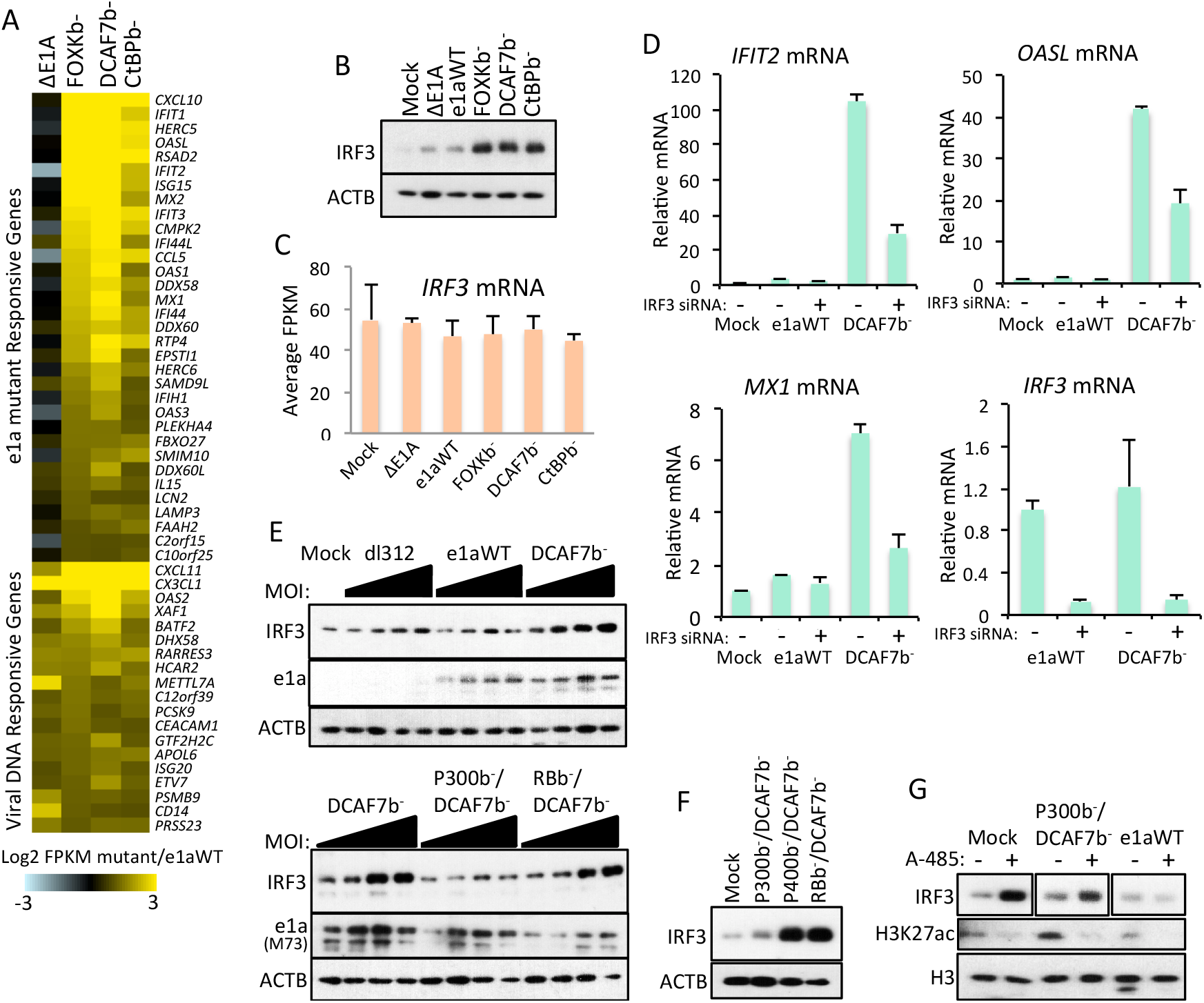
e1a inhibition of p300/CBP lysine-acetyl transferase activity induces IRF3 stabilization. (A) Heatmap displaying changes in RNA-seq FPKM values of the 52 host genes expressed at >2-fold higher level in primary HBTECs following infection by vectors for e1a C-terminal mutants compared to a vector for wt e1a. Cluster 1 = e1a C-terminal mutant responsive genes activated <2-fold by the introduction of viral DNA during infection since they were not induced by infection with *dl312* (ΔE1A, (*21*)), which undergoes a normal infection process but does not express viral proteins. Cluster 2 genes were activated >2-fold by the introduction of viral DNA during infection, or some other aspect of the adenovirus infection process, since they were induced >2-fold by infection with *dl*312 (ΔE1A). (B) Western blots of protein from 24h p.i. of HBTECs with Ad5 vectors expressing wt e1a or the e1a mutants indicated at the top. (C) qRT-PCR with RNA extracted from HBTECs 24h p.i. with Ad5 vectors expressing wt e1a or e1a mutants indicated at the top. (D) qRT-PCR with RNA extracted from HBTECs transfected for two days with the indicated siRNAs, and infected with Ad5 vectors expressing the e1a mutants indicated at the top for an additional 24h. (E) Western blots with protein from 24h p.i. of HBTECs infected with Ad5 vectors expressing the indicated e1a mutant at increasing MOI: 20, 60, 100, 140. (F) Western blots of protein from HBTECs infected with Ad5 vectors expressing the indicated e1a mutants. (G) Western blots of protein from 12h-infected + 12h DMSO or 10 µM A485-treated HBTECs. (C, D) Data represent averages of three experimental replicates + SD. (D, G) “-” indicates negative control siRNA (siCtrl).

The increase in ISG expression following infection with each of the e1a C-terminal mutants coincided with an increase in IRF3 protein (Figure 1B), without an increase in IRF3 mRNA (Figure 1C). Pulse-chase radio-labeling analysis with ^35^S-Met/Cys showed that this increase in IRF3 was due to an increase in IRF3 protein stability (*18*). A siRNA knockdown (KD) of IRF3 in HBTECs significantly reduced the ability of DCAF7b-e1a to activate *IFIT2, OASL*, or *MX1* (Figure 1D). Therefore, e1a C-terminal mutants trigger a late interferon response (*28*) in infected HBTECs by stabilizing IRF3 by a mechanism that is not triggered by viral DNA alone, but requires binding of FOXK1 or 2, the DCAF7-protein kinase complex, and CtBP1 or 2 to the same e1a molecule (*18*).

### Inhibition of p300/CBP acetyl transferase activity triggers IRF3 stabilization

Most of e1a’s ability to regulate transcription is credited to interactions of the N-terminal half of e1a with p300/CBP and RB family proteins (RBs) (Figure S1) (*10*). Interactions between the N-terminal E1A region and chromatin remodeling/histone acetyl transferase complexes containing p400 and/or TRRAP also have been described (Figure S1) (*17*). To determine if these interactions with the N-terminal half of e1a are necessary for IRF3 stabilization in response to e1a C-terminal mutants, we constructed three Ad5 vectors that express double-interaction mutants of e1a: e1a defective in binding both p300/CBP and DCAF7 (P300b-/DCAF7b-), e1a defective in binding both RBs and DCAF7 (RBb-/DCAF7b-), or e1a defective in binding both p400/TRRAP and DCAF7 (p400b-/DCAF7b-). The e1a mutations eliminating binding to p300/CBP or RBs contained two or four amino acid changes from the wt sequence and are described in our previous report (*10*), and e1a binding to p400 and TRRAP was eliminated by deletion of aa 26-35 (*17*). Infection of HBTECs with increasing moi (20 – 140) of *dl*312 (ΔE1A) or the vector expressing wild-type small e1a (e1aWT) affected IRF3 level only modestly (Figure 1E upper). But, as before (Figure 1B), infection with the Ad vector for the DCAF7b-e1a mutant greatly increased the level of IRF3 (Figure 1E, upper). RBb-/DCAF7b-e1a also induced IRF3 accumulation. However, the P300b-/DCAF7b-e1a mutant induced much less accumulation of IRF3 (Figures 1E lower, 1F). Mutant p400b-/DCAF7b-retained the ability to induce IRF3 accumulation similarly to the RBb-/DCAF7b-e1a mutant (Figure 1F). Therefore, e1a C-terminal mutants must bind p300/CBP to induce IRF3 stabilization, but not RBs, p400, or TRRAP.

Small e1a can inhibit p300/CBP histone acetylase activity by binding p300/CBP through the e1a N-terminal ∼15 amino acids plus the C-terminal portion of CR1 (amino acids 54-71) (*14*) (Figure S1). To determine if e1a inhibition of p300/CBP lysine-acetyl transferase activity is responsible for increasing IRF3 stability in cells infected with the e1a C-terminal mutants, we performed experiments with A-485, a potent and specific small molecule inhibitor of p300 and CBP acetyl transferase activities (*29*). p300/CBP-mediated acetylation of histone H3 lysine 27 (H3K27ac) was greatly reduced by A-485 treatment for 12 h (Figure 1G). In contrast, A-485 treatment increased IRF3 protein in both mock- and P300b-/DCAF7b-vector-infected HBTECs, but not in HBTECs infected with the vector for wt e1a (Figure 1G). These data show that inhibition of p300/CBP acetyl transferase activity by either A-485 or the N-terminal region of e1a induces IRF3 stabilization. Since wt e1a inhibits p300/CBP but does not increase IRF3 stability, we hypothesized that wt e1a prevents IRF3 stabilization, and that this activity of wt e1a is lost when any one of the e1a interactions with host proteins made by the three C-terminal e1a conserved regions fails to occur.

### Interactomes of wt e1a and e1a C-terminal mutants

To investigate why wt e1a, but not the e1a C-terminal mutants, prevents IRF3 stabilization following infection, we characterized wt e1a-binding proteins and proteins bound by each of the e1a C-terminal mutants. We performed immunoprecipitation (IP) of e1a from lysates of A549 cells (a human bronchial carcinoma cell line) after mock-infection, or infection with Ad vectors expressing wt e1a, or the FOXKb-, DCAF7b-, or CtBPb-e1a mutants. Immunoprecipitation was performed using a monoclonal antibody (M58) that binds in e1a’s N-terminal half (*30*). This was followed by mass spec to identify proteins that were differentially bound to wt e1a compared to the three e1a C-terminal mutants (Table S1).

Specific disruption of FOXK TF-binding to the FOXKb-e1a mutant, DCAF7 and DYRK1A/B binding to the DCAF7b-e1a mutant, and CtBP1 or 2 binding to the CtBPb-e1a mutants, observed earlier by IP-westerns (*18*), were validated by these mass spec results (Figure 2A). Several known e1a binding proteins that bind peptides in the N-terminal half of e1a were greatly enriched in the anti-e1a IPs with wt e1a and e1a C-terminal mutants compared to mock-infected cells, confirming that wt and mutant e1a-IPs were effective and specific. Among the proteins co-IPed with wt and C-terminal mutant e1as, CBP, p300, RB, RBL2, RBL1, TRRAP, and p400 had the highest enrichments (Figure 2A). Other previously reported e1a interacting proteins were also detected such as MYC (*31*) and 26S proteasome subunits (*32*). 191 identified proteins were >5-fold enriched over mock for binding to wt e1a and at least two of the three e1a C-terminal mutants (to account for disrupted binding to one of the e1a C-terminal mutants). These 191 proteins were significantly enriched for known protein-protein interactions compared to random proteins (p < 1×10^−6^, Figure 2B). Enriched KEGG pathways gene ontologies for e1a bound proteins included cell cycle, pathways in cancer, TGFβ signaling, mismatch repair, glycolysis, viral carcinogenesis, proteasome, and spliceosome (Figure 2B). Among the previously unreported e1a bound proteins was PRKDC, the catalytic subunit of DNA-PK, a complex involved in double-stranded DNA break repair, which was recently reported to be antagonized by e1a (*33*). We also observed binding of Ribosomal Protein Kinase S6 A4 (KS6A4) to wt, FOXKb-, and CtBPb-e1as but not to DCAF7b-e1a (Figure 2A), suggesting that KS6A4 is another protein kinase in addition to DYRK1A, DYRK1B and HIPK2 in protein complexes with e1a and DCAF7 (*34*). Additional key proteins of signaling pathways that regulate cell growth were found to co-immunoprecipitate with e1a including the MAPK signaling pathway kinase ARAF, TGFβ-signaling pathway-regulated transcription factor SMAD3, and epidermal growth factor receptor EGFR (Figure 2B).

**Figure 2.**
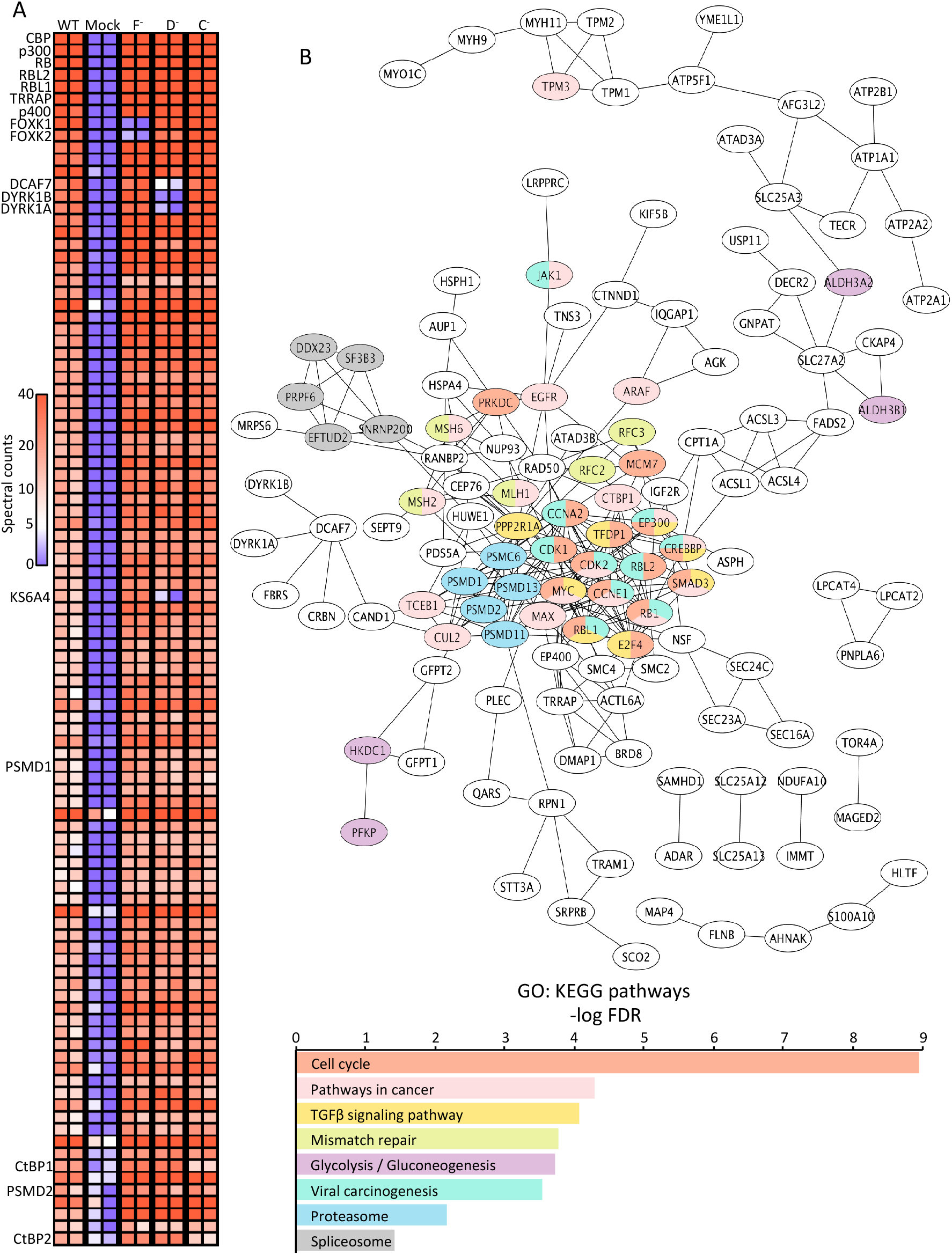
e1a interactome. (A) Heatmap displaying values for spectral counts of peptides from proteins detected by mass spec from M58 (anti-e1a) IPs. Values for duplicate mass spec runs are represented in adjacent columns. Heatmap is limited to proteins with ≥ an average of 10 spectral counts between duplicates and ≥5-fold enrichment of wt e1a over Mock. (B) Top: Cytoscape protein-protein interaction map for 191 proteins that were enriched for binding to wt e1a and at least two e1a C-terminal mutants > 5-fold over mock. Colors refer to the gene ontology shown below. Bottom: Select enriched KEGG Pathways gene ontologies for proteins represented in Cytoscape protein-protein interaction network.

### DCAF10 associates with and destabilizes e1a and regulates IRF3 stability

The most differentially bound protein between wt e1a and the three e1a C-terminal mutants was DCAF10, which bound to wt e1a but none of the e1a C-terminal mutants in both A549 and HBTECs (Figure 3A). DCAF10 (aka WDR32) was identified as a DDB1-interacting WD-repeat-containing protein that may act as a substrate receptor for CUL4-DDB1 E3 ubiquitin ligase complexes (i.e., a Cullin 4-ring ligase or “CRL4”) (*35*). The specific interaction of DCAF10 with wt e1a but not e1a C-terminal mutants was confirmed by e1a IP-western blot from infected A549 cells expressing HA-tagged DCAF10 (Figure 3B). KD of DCAF10 in HBTECs, to ∼30-40% the mRNA level in negative control (siCtrl) transfected cells (Figure 3C), increased wt e1a protein, but did not further increase the high level of the e1a C-terminal mutants that do not associate with DCAF10 (Figure 3D). While DCAF10 KD caused wt e1a protein to increase, it had no effect on e1a mRNA levels (Figure 3E). These results suggest that DCAF10 binding to e1a (directly or indirectly) destabilizes e1a. Treatment with MLN4924, a selective inhibitor of NEDD8-activating enzyme required for activation of Cullin-based E3 ubiquitin ligases (*36*), confirmed that e1a degradation depends on ubiquitinylation by a Cullin-based E3 (Figure 3F).

**Figure 3.**
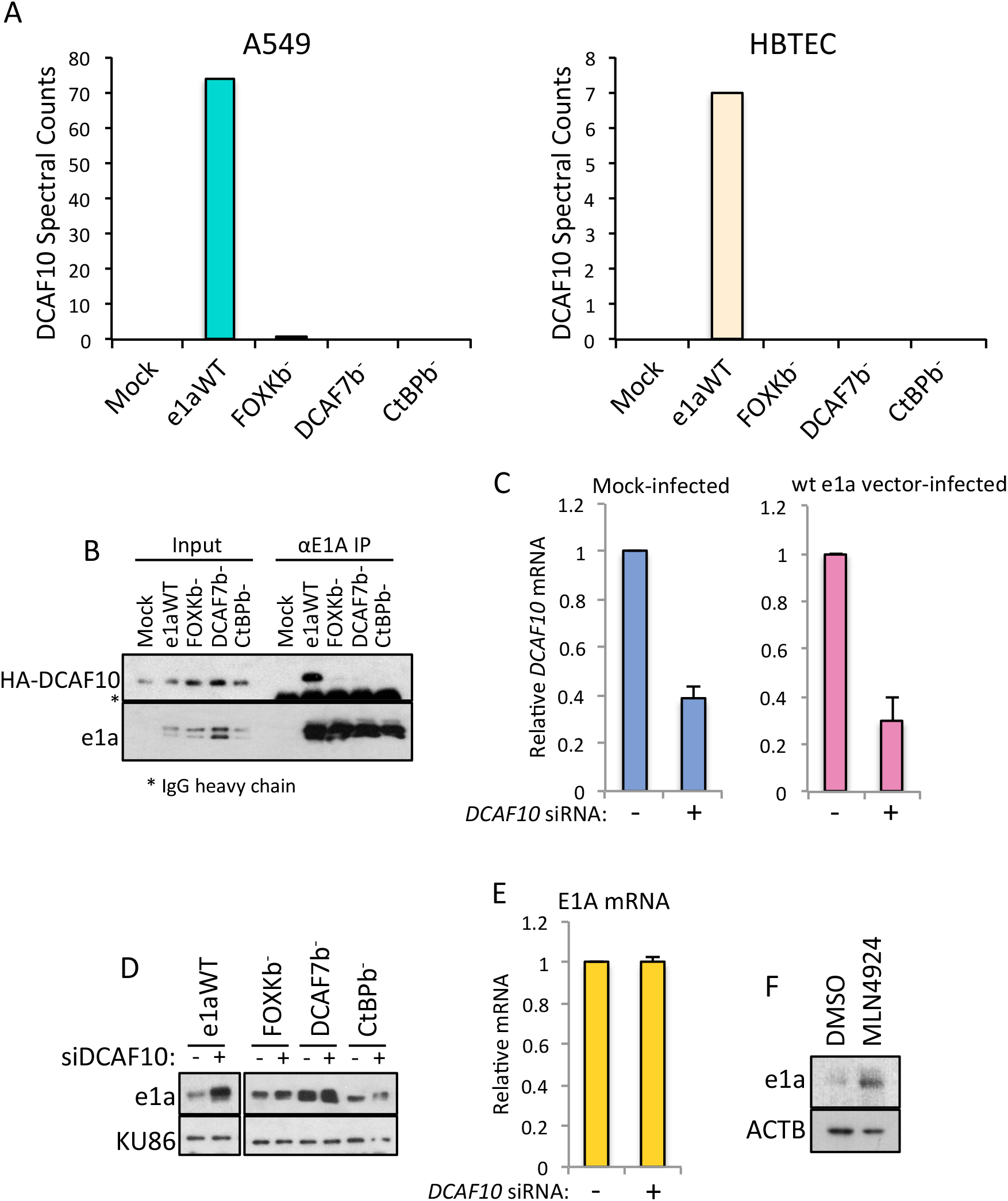
Wild-type e1a binds DCAF10 but not e1a C-terminal mutants. (A) Spectral counts for DCAF10 mapped peptides from anti-e1a (M58) IP mass spec from HBTEC and A549 24h p.i. with vectors for wt or C-terminal mutant e1a as indicated at the bottom. (Figure 3 legend continued). (B) Western blots following M58 (anti-e1a) IP of protein from A549 24h p.i. with vectors for wt e1a or the indicated e1a C-terminal mutants. (C) qRT-PCR with cDNA prepared from HBTECs transfected with control or anti-siRNA for two days, and then mock-infected (Left) or infected with the wt e1a vector (Right). (D) Western blots with protein from HBTECs transfected for two days with control or anti-DCAF10 siRNA, and then infected for 24h with the vectors for wt or C-terminal mutant e1as indicated at the top. (E) qRT-PCR for E1A mRNA with cDNA prepared from HBTECs transfected for two days with control (-) or DCAF10 siRNA (+), and then infected with the e1a wt vector. (F) Western blots with protein from HBTECs infected with e1a wt vector for 24h, and then treated for 6h with control DMSO or 20μM MLN4924. (C,D,E) “–” indicates negative control siRNA (siCtrl). (C,E) Data represent averages of three experimental replicates + SD.

To determine if DCAF10 regulates IRF3 stability, we performed DCAF10 KD in mock-infected HBTECs and HBTECs infected with the vector for wt e1a. Western blotting for e1a and IRF3 showed that KD of DCAF10 caused an increase in IRF3 protein in both mock-infected HBTECs and HBTECs expressing wt e1a (Figure 4A), without increasing IRF3 mRNA (Figure 4B). DCAF10 KD also activated *OASL* and *IFIT2* mRNA expression, ISGs induced by the e1a C-terminal mutants, in uninfected HBTECs and HBTECs expressing wt e1a (Figure 4C), indicating that the increased IRF3 protein was functional for activating ISG transcription. These data suggest that DCAF10 inhibits ISG activation by the e1a C-terminal mutants because it destabilizes IRF3. The data also suggest that e1a-association with DCAF10 enhances DCAF10’s ability to destabilize IRF3, preventing IRF3 accumulation and activation of ISGs as a consequence of e1a inhibition of p300/CBP acetyl transferase activity.

**Figure 4.**
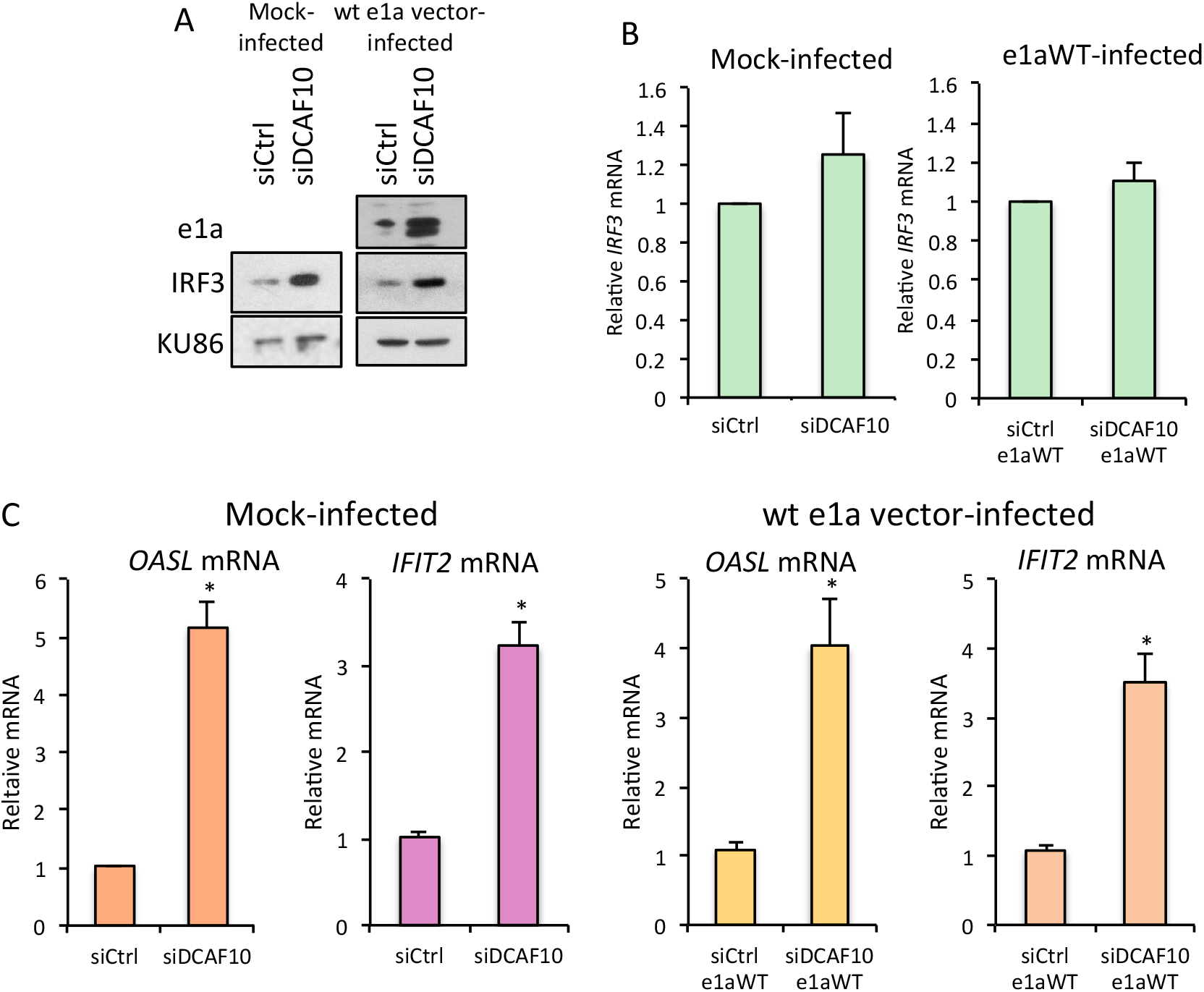
DCAF10 destabilizes IRF3 and inhibits ISG activation. (A) Western blots of protein from HBTECs transfected with control or DCAF10 targeting siRNAs for two days, then mock- or infected with the Ad5 vector that expresses wt e1a for 24h. (B) qRT-PCR with RNA extracted from HBTECs transfected with control or DCAF10 siRNA for two days, and then 24h mock-infected (Left) or infected with the Ad5 vector expressing e1a wt (Right). (C) qRT-PCR with RNA extracted from HBTECs transfected for two days with the indicated siRNAs and then mock-infected (Left two) or infected with the Ad5 vector expressing wt e1a (Right two) for 24h. (B, C) Data represent averages of three experimental replicates + SD. “*” indicates p<0.01.

### DCAF10 association with e1a promotes assembly of a CRL4 E3 ubiquitin ligase

To investigate the mechanism of DCAF10 regulation of IRF3 stability we searched for proteins bound by DCAF10 in A549 and HBTECs by expressing HA-tagged DCAF10 from an Ad vector that does not express E1A proteins. We used an Ad vector in order to express HA-tagged DCAF10 in nearly one-hundred percent of the primary HBTECs as well as A549 cells. Proteins associated with HA-DCAF10 were identified by mass spec of anti-HA IPs prepared under conditions that maintain specific protein-protein interactions. We observed co-immunoprecipitation (co-IP) of DDB1, CUL4A, and CUL4B with HA-DCAF10 when wt e1a was co-expressed (Figures 5A, B), supporting the hypothesis that e1a forms a CUL4-based E3 ubiquitin ligase complex with DCAF10. Of considerable significance, although DDB1 co-IPed with HA-DCAF10 in the absence and presence of wt e1a, the critical subunits CUL4A or B required for E3 ubiquitin ligase activity, did not co-IP with DCAF10 in the absence of wt e1a (Figure 5B). This result indicates that wt e1a promotes assembly of an e1a-DCAF10-containing Cullin 4-based E3 ubiquitin ligase complex (i.e., a Cullin4-Ring E3 Ligase, or “CRL4”). Consistent with this, wt e1a bound more CUL4A/B than the e1a C-terminal mutants that do not associate with DCAF10 (Figure S2). Earlier gel filtration analysis of nuclear extract from 293 cells that express E1A proteins showed that the major small e1a-containing complex(es) was ∼1 MDa (*10*), large enough to include e1a, Cullin 4A or 4B, DDB1, FOXK1 or 2, CtBP1 or 2, and DCAF7 and its associated protein kinases (*12*).

**Figure 5.**
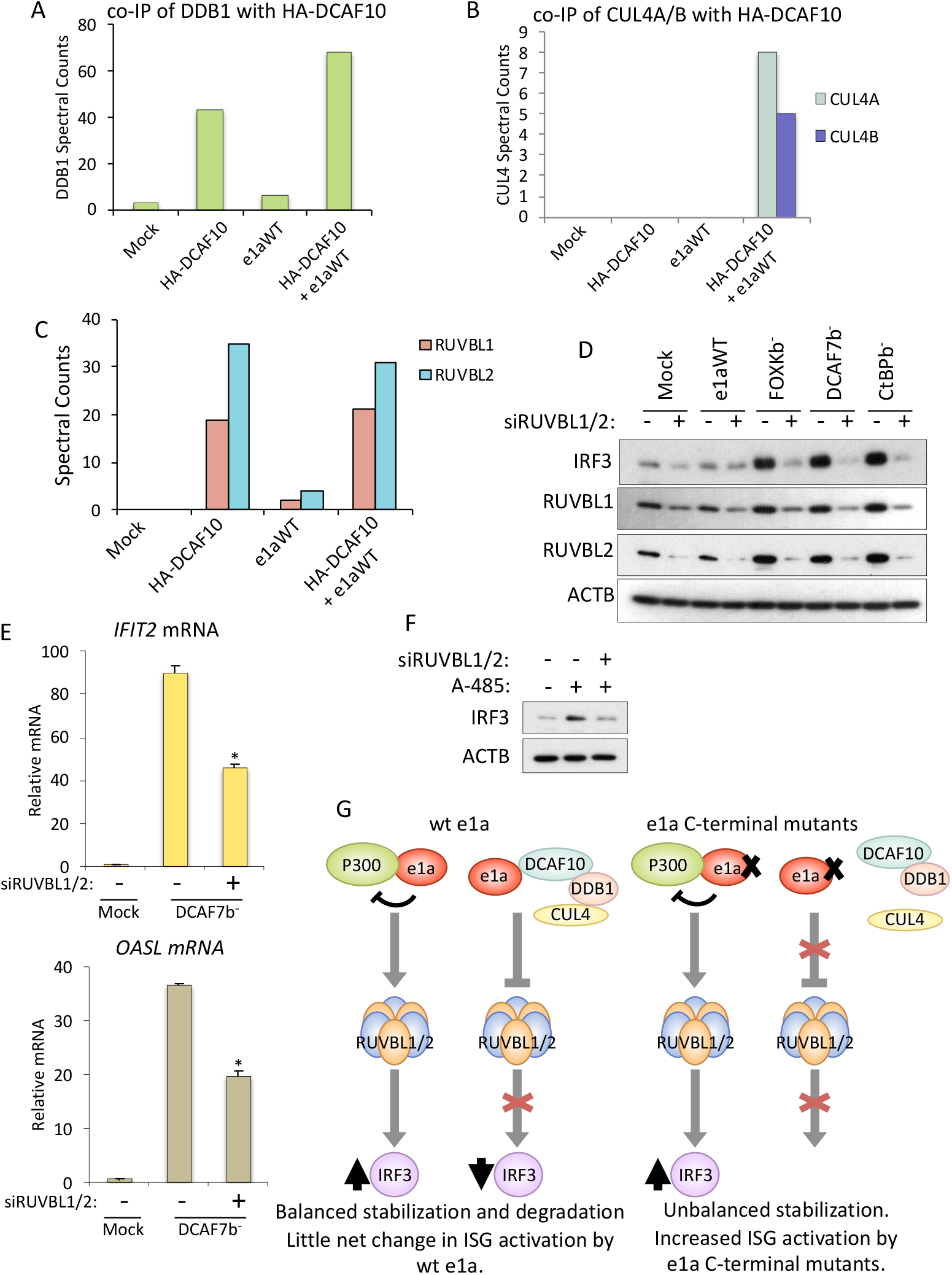
e1a C-terminal mutant stabilization of IRF3 requires RUVBL1/2. (A) DDB1, (B) RUVBL1/2, (C) CUL4A/B Spectral counts in anti-HA IPs from cells expressing HA-DCAF10 for. (D) Western blots for the proteins indicated at the left from HBTECs treated for two days with control siRNA (-) or siRNAs for RUVBL1 and 2 (+), and then mock-infected or infected with the Ad5 vectors indicated at the top for 24h. (E) qRT-PCR of IFIT2 and OASL mRNA from HBTECs treated as in (D.) Averages of three experimental replicates + SD. “*” indicates p<0.01 for the difference between control siRNA and RUVBL1/2 siRNA-transfected HBTECs. (F) Western blots for IRF3 or ACTB from HBTECs transfected with control siRNA (-) or siRNAs for RUVBL1/2 (+) for 2 days then treated for 12h with DMSO or 10µM A485. (G) A model of e1a inhibition of P300 causing stabilization of IRF3 mediated through a RUVBL1/2-containing complex, and a complex of e1a-DCAF10-DDB1-CUL4 that antagonizes IRF3 stabilization.

Importantly, we did not detect co-IP of IRF3 with HA-DCAF10. This argues against a direct interaction between the two proteins, and, therefore, against direct ubiquitinylation of IRF3 by the e1a-DCAF10-CRL4. We did find that HUWE1, a HECT-type E3 ubiquitin ligase also targeted by the HIV Vpr-CLR1 E3 (*29, 40*), co-IPed specifically with HA-DCAF10, with reduced co-IPed HUWE1 after expression of e1a (Figure S3). This reduced HUWE1 in the presence of e1a may have been due to enhanced polyubiquitinylation and proteasomal degradation of HUWE1 in the presence of e1a. However, HUWE1 did not appear to be directly involved in regulating IRF3 stability, because HUWE1 KD did not alter IRF3 levels in uninfected or e1a C-terminal mutant-expressing HBTECs (data not shown).

### AAA+ ATPases RUVBL1 and 2 are targets of the e1a-DCAF10-CRL4

The AAA+ family ATPases RUVBL1 and RUVBL2 (a.k.a. Pontin and Reptin, respectively) that form complexes involved in DNA damage sensing and repair (*38*) and were previously reported to be required for maximal transcriptional responses to interferon α (*26, 27*) were also found in HA-DCAF10 IPs, whether or not e1a was co-expressed (Figure 5C). However, unlike HUWE1 KD, double KD of both RUVBL1 and 2 affected IRF3 level dramatically, preventing IRF3 protein accumulation in HBTECs expressing e1a-C-terminal mutants, and also reducing IRF3 protein in mock-infected HBTECs (Figure 5D), with only modest changes in IRF3 mRNA levels (Figure S4). These results demonstrate that RUVBL1/2 are necessary for stabilization of IRF3 in HBTECs expressing e1a C-terminal mutants. RUVBL1 and RUVBL2 proteins also increased after expression of the e1a C-terminal mutants (Figure 5D), without increases in their mRNAs (Figure S5), suggesting that DCAF10 regulates the stabilities of RUVBL1 and 2 as well as IRF3. Importantly for the mechanism of e1a inhibition of innate immunity, RUVBL1/2 KD inhibited activation of ISGs by the e1a C-terminal mutants (Figure 5E), probably because KD of RUVBL1/2 inhibited stabilization of IRF3 by the e1a C-terminal mutants (Figure 5D).

The results above indicate that RUVBL1/2 are required for e1a C-terminal mutants to stabilize IRF3. To determine if RUVBL1/2 also are necessary for the IRF3 stabilization observed with inhibition of p300/CBP acetyl transferase activity (Figure 1G), we treated HBTECs with A-485 with or without RUVBL1/2 KD (Figure 5F). A-485 treatment increased IRF3 protein as before, but RUVBL1/2 KD prevented the A-485-induced increase (Figure 5F). Taken together these data suggest that inhibition of p300/CBP acetyl transferase activity by A-485, or by the e1a C-terminal mutants, leads to stabilization of IRF3 dependent on RUVBL1/2 activity, and that the e1a-DCAF10 interaction inhibits RUVBL1/2 from stabilizing IRF3 and activating interferon-stimulated genes (Figure 5G).

Since our results were consistent with the model that e1a assembles a DCAF10-containing CRL4 E3 ubiquitin ligase complex, we searched for proteins that might be substrates for this complex in the e1a-IPs. HUWE1 and DNAJA1 co-IPed with both e1a and HA-DCAF10 (Figure 6A, B), raising the question of whether they might be substrates for the e1a-containing E3 ubiquitin ligase complex. Consistent with this, western blots for HUWE1 and DNAJA1 from infected HBTECs showed that both proteins had higher levels when e1a C-terminal mutants were expressed compared to wt e1a (Figure 6C), although their mRNA levels were not higher (Figure 6D). Additionally, more HUWE1 and DNAJA1 associated with e1a C-terminal mutants compared with wt e1a (Figure 6B). This could be due to a lower rate of HUWE1 and DNAJA1 degradation when e1a cannot complex with DCAF10. These results are consistent with the model that e1a redirects a DCAF10-CRL4 to target e1a-bound proteins such as HUWE1 and DNAJA1, as well as RUVBL1 and 2 and other potential substrates, for ubiquitin ligase-mediated proteasomal degradation.

**Figure 6.**
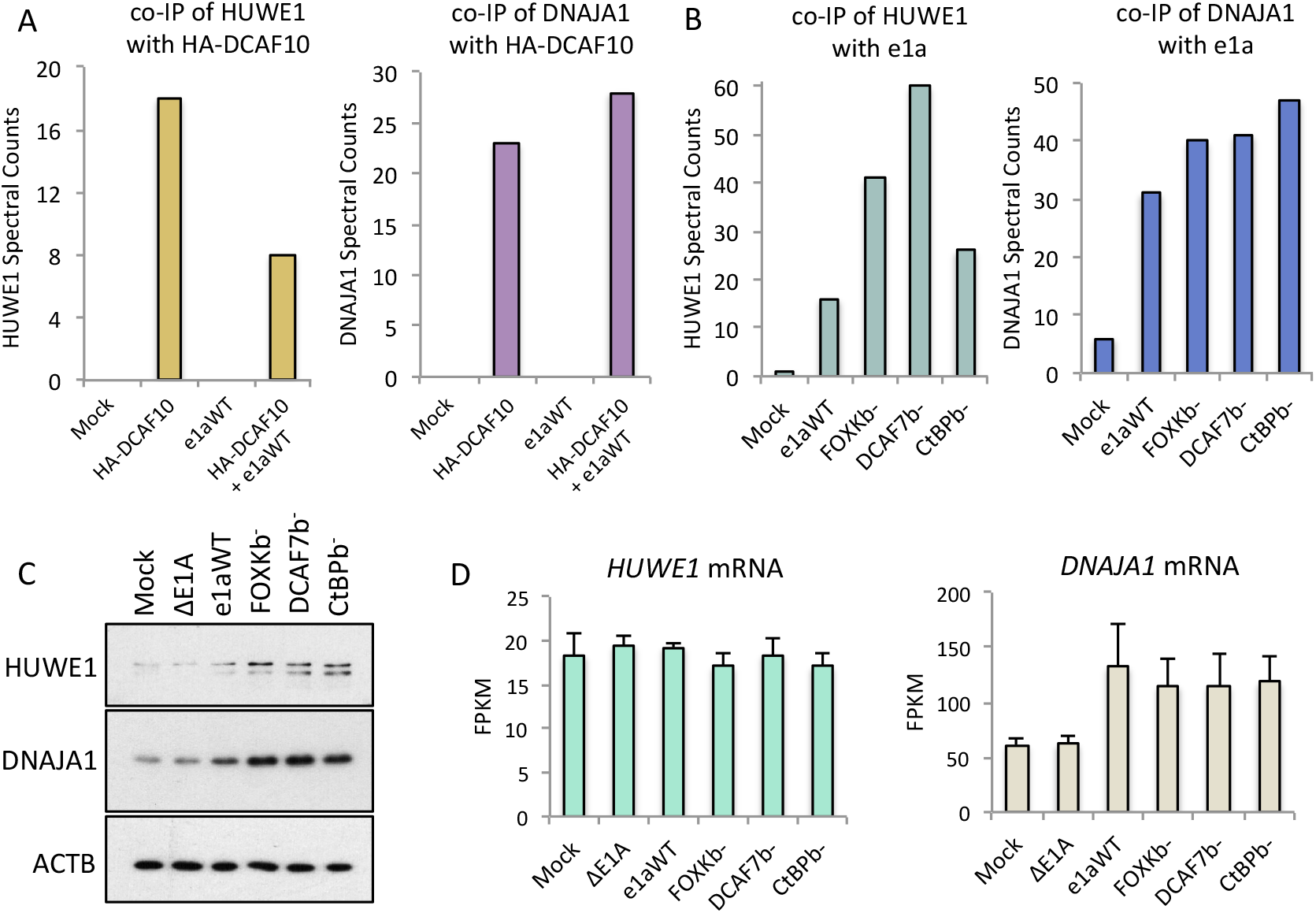
e1a bound proteins HUWE1 and DNAJA1 accumulate when e1a cannot bind DCAF10. (A) Spectral counts for HUWE1 (left) and DNAJA1 (right) peptides from HA-DCAF10 (anti-HA) IP mass spec from A549 cells 24h p.i. (B) Spectral counts for HUWE1 (left) and DNAJA1 (right) peptides from M58 (anti-e1a) IP mass spec from A549 cells 24h p.i. (C) Western blots with protein from HBTECs 24h p.i. with the indicated Ad5 vectors. (D) FPKM of HUWE1 (left) and DNAJA1 (right) from RNA-seq of 24h-infected HBTECs. Data represent averages of three experimental replicates + SD.

## DISCUSSION

We identified an activity of adenovirus e1a that triggers an anti-viral innate immune response, and an additional e1a activity that counteracts this host anti-viral response. Previously, we reported that adenovirus e1a C-terminal mutants defective for binding cellular proteins FOXK1/2, DCAF7, or CtBP1/2, potently activate ∼50 interferon-stimulated genes (*18*). Activation of these ISGs required IRF3, which was stabilized (*18*) and accumulated to high concentration when any one of these e1a mutants was expressed (Figure 1B). By 12 hours post infection with Ad vectors for these e1a mutants, IRF3 was transported into nuclei and became chromatin bound before it was phosphorylated at its previously discovered activating site S396 (*39*). These observations led us to investigate how expression of e1a mutant proteins, and not adenovirus nucleic acids, triggered this anti-viral response, and how the response was prevented by wt e1a dependent on the three e1a C-terminal interactions with FOXK1/2, DCAF7, and CtBP1/2. The observations that infection with the E1A deletion mutant *dl*312 does not cause stabilization of IRF3 or activation of multiple ISGs indicates that adenovirus DNA alone does not activate the innate immune response in adenovirus-infected cells. This may be because adenovirus DNA is sterically blocked from detection by TLR9 in endosomes and by cyclic GMP-AMP synthase (cGAS) and other sensors for DNA in the cytoplasm (*25*) because it is covered by virion protein VII (*23*)).

### Inhibition of p300 and CBP lysine-acetyl transferase activity leads to IRF3 stabilization and activation of ISGs

Ad5 vectors expressing e1a double interaction mutants defective for binding p300/CBP and DCAF7, or RBs and DCAF7, or p400/TRRAP and DCAF7 revealed that e1a C-terminal mutants must bind p300/CBP through their N-terminal regions to trigger IRF3 stabilization (Figure 1E, F). Also, inhibition of p300/CBP lysine-acetyl transferase (KAT) activity with the highly specific small molecule inhibitor A-485 (*29*) was sufficient to induce IRF3 stabilization (Figure 1G), revealing that inhibition of p300/CBP KAT enzymatic activity is responsible for triggering this innate immune response. Since p300/CBP are common targets of DNA tumor viruses, e.g. HPV E6 (*40*), mouse polyomavirus and SV40 large T antigens (*41, 42*), and adenovirus E1As (*43*–*45*), IRF3 stabilization in response to p300/CBP inhibition may have evolved as an anti-viral host defense. Since this previously uncharacterized activation of an innate immune response by inhibition of p300/CBP KAT activity was revealed by e1a mutants defective for any one of three e1a C-terminal protein interactions, and wt e1a prevented IRF3 stabilization, we hypothesize that wt e1a evolved a separate activity dependent on the three C-terminal interactions in order to retain p300/CBP inhibition, but counter the resulting IRF3 stabilization that otherwise induces ISGs and inhibits viral replication (*18*).

Our earlier complementation analysis indicated that a single e1a protein molecule must bind FOXK1 or 2, the DCAF7-complex, and CtBP1 or 2 to prevent activation of ISG expression (*18*). These results suggest that assembly of a complex containing e1a and these three seemingly unrelated host proteins prevents IRF3 stabilization that is otherwise induced by e1a inhibition of p300/CBP acetyl transferase activity.

### DCAF10 associates with e1a dependent on all three e1a C-terminal interactions with seemingly unrelated host cell proteins

To explore why all three of the e1a C-terminal mutants fail to prevent IRF3 stabilization, we searched for proteins differentially bound to wt e1a and all three of the e1a C-terminal mutants. DCAF10 fulfilled these criteria: it associated with wt e1a but none of the three e1a C-terminal mutants (Figures 3A, B). A possible explanation for why three different, well-separated mutations in e1a’s C-terminal half resulted in loss of DCAF10 binding may be because of common alterations in e1a phosphorylation of all three mutants compared to wt e1a. Previously, we reported that wt e1a is phosphorylated at one or more sites that cause a mobility shift during SDS-PAGE, which is (are) not phosphorylated on the e1a C-terminal mutants (*18*). Phosphorylation of this site(s) may be necessary for DCAF10 binding to e1a, making the site a “phospho-degron,” a strategy commonly used to regulate protein polyubiquitinylation and proteasomal degradation. We also found that DCAF10 association destabilized wt e1a (Figure 3D). This is consistent with the previous identification of a phospho-site between e1a amino acids 178-192 that decreases e1a stability (*46*). Together with our previous report, this lack of a mobility shift for all three C-terminal mutants suggests that e1a phosphorylation at a specific site(s) is required for DCAF10-binding to e1a, and that e1a polyubiquitinylation and proteasomal degradation is decreased for all three of the e1a C-terminal mutants because they fail to be phosphorylated at this site(s). One or more of the DCAF7 associated protein kinases DYRK1A, DYRK1B, HIPK2 (*34*) or KS6A4 (Figure 2A) may phosphorylate this site(s) in e1a, stimulating binding of DCAF10 and assembly of the complete e1a-DCAF10-CRL4.

### e1a-DCAF10 targeting of RUVBL1/2 prevents IRF3 stabilization and activation of ISGs

We did not detect a direct interaction between DCAF10 and IRF3 in the presence or absence of e1a, suggesting that IRF3 is not directly polyubiquitinylated by an e1a-DCAF10-CRL4 E3. However, we did detect association of epitope-tagged DCAF10 with RUVBL1 and 2 (Figure 5C). Stabilization of RUVBL1/2 by the e1a C-terminal mutants also was observed, and RUVBL1/2 were required for stabilization of IRF3 by the e1a C-terminal mutants (Figure 5D). Since the e1a C-terminal mutants caused an increase in RUVBL1/2 proteins but not their mRNAs (Figures 5D, S5), they likely cause stabilization of RUVBL1 and 2 protein. In contrast, wt e1a prevented RUVBL1/2 stabilization (Figure 5D). We proposed that this is because wt e1a forms the complete CRL4 complex that directs polyubiquitinylation and proteasomal degradation of RUVBL1 and 2. RUVBL1 has been reported to co-immunoprecipitate with the C-terminal half of e1a (*27*). This might be due to an indirect interaction between e1a and RUVBL1 mediated by DCAF10.

Detection of higher levels of DDB1 and CUL4A and B binding to HA-DCAF10 in the presence of e1a (Figures 5A, B) suggests that e1a promotes DCAF10 incorporation into this CRL4 complex. Also, the stabilization of wt e1a by knocking-down DCAF10, which is not observed for the e1a C-terminal mutants unable to bind DCAF10 (Figure 3D), suggests that DCAF10 binds to wt e1a as part of an active ubiquitin ligase complex that polyubiquitinylates wt e1a as well as other proteins bound to wt e1a including RUVBL1 and 2. Our observations that e1a bound proteins HUWE1 and DNAJA1 accumulated to higher levels when e1a had mutations blocking the e1a-interaction with DCAF10 (Figure 6C) are consistent with this model that e1a directs substrate targeting by this DCAF10-containing CRL4 ubiquitin ligase to proteins bound by e1a. A similar activity has been well-documented for HIV-1 Vpr, which binds DCAF1 and forms a DDB1-CRL4 E3 ubiquitin ligase that polyubiquitinylates several cellular proteins that inhibit HIV replication, such as uracil DNA glycosylase UNG2 (*47*).

A surprising observation was that RUVBL1/2 knock-down prevented stabilization of IRF3 by the e1a C-terminal mutants and by inhibition of p300/CBP KAT activity with A-485 (Figures 5D, F). IRF3 stabilization was presumably due to inhibition of IRF3 polyubiquitinylation following expression of the e1a mutants or treatment with A-485. Also, Olanubi et al. (*27*) reported that the e1a C-terminal interaction with RUVBL1 observed by co-immunoprecipitation with E1A is required for E1A to repress activation of ISGs by IFNα. These observations might be explained if protein conformational changes regulated by RUVBL1/2 were required for IRF3 polyubiquitinylation. For example, RUVBL1/2 might be required subunits of the cellular E3(s) for IRF3; or might be required for signaling to a cellular IRF3 E3 in response to p300/CBP-inhibition; or may be necessary to push IRF3 into a conformation that reveals a binding site for an IRF3 E3, among other possibilities.

RUVBL1/2 function as co-chaperones for HSP90 and are involved in diverse cellular processes as components of several different multiprotein complexes (*48, 49*). In the cytoplasm, RUVBL1/2-containing complexes function in the assembly of several essential multi-molecular complexes such as snRNPs, snoRNPs, the telomerase RNP complex, RNA Pol2, the TRAPP chromatin remodeling complex, and the PIKK family of multi-subunit protein kinases including mTORC1, which regulates ribosome and protein synthesis, and SMG-1 required for nonsense-mediated mRNA decay (NMD). In the nucleus, RUVBL1/2 regulate transcription and DNA double-strand break repair as subunits of chromatin remodeling complexes TIP60 and INO80. They also associate with and regulate the Fanconi Anemia DNA repair complex (*50*) and MYC (*51*), which regulates protein synthesis and cell size (*52*). We speculate that the connection between RUVBL1/2 and IRF3 stabilization is mediated by the chaperone-like complex R2TP, consisting of RUVBL1, RUVBL2, PIH1D1, and RPAP3 (*53*). All four of these host proteins were enriched in anti-HA IPs of extracts from cells expressing HA-DCAF10 (Table S2). R2TP activity is required to maintain the stability of these several RUVBL1/2-containing multi-protein complexes (*50, 54*).

AAA+ ATPases such as RUVBL1/2 are the “moving parts” of several cellular multi-protein machines. They undergo large conformational changes when ATP is exchanged for ADP in their nucleotide binding pocket, when that ATP is hydrolyzed to ADP and phosphate, and when the phosphate is released, preparing the protein machine for another cycle of large conformational changes, as in the case of the myosins. Such large, regulated changes in protein conformation may be required to expose sites in RUVBL1/2 and IRF3 to ubiquitin ligases. Whatever the molecular mechanism, it is clear from the results reported here that RUVBL1/2 are required to activate innate immunity in response to inhibition of p300/CBP KAT activity. It will be fascinating to pursue additional studies that explain why the RUVBL1/2 AAA+ ATPases are required for IRF3 stabilization when p300/CBP KAT activity is inhibited.

## Materials and Methods

### Cell Culture

Cells were maintained at 37°C in a 5% CO_2_ incubator. A549 cells were grown in Dulbecco’s Modified Eagle Medium (DMEM) with 10% fetal bovine serum. Human Bronchial/Tracheal Epithelial Cells (HBTEC, Lifeline Cell Technology Cat# FC-0035; lot# 02196) in BronchiaLife Medium Complete Kit (Lifeline Cell Technology catalog number: LL-0023).

### Ad Vectors and Infection

Ad5 vectors expressed Ad2 wt or mutant e1a’s from the normal E1A promoter with the dl1500 deletion removing the 13S E1A mRNA 5’ splice site (*2*). The vectors were constructed using the Ψ5 vector and in vivo Cre-mediated recombination (*55*), and consequently contain an out of frame insertion of a LoxP site at the Bgl II site in the region encoding the carboxy-terminus of E1B-55K. All infections were for 24 h at an moi of 60, unless otherwise indicated. The ΔE1A mutant was dl312 (*21*).

Additional Materials and Methods can be found in the Supplementary Materials

## Supplementary Materials

Supplementary Materials and Methods

Figure S1. Host proteins that bind E1A regions in the primate adenoviruses.

Figure S2. e1a C-terminal mutants have reduced binding to CUL4A/B.

Figure S3. DCAF10 binds HUWE1.

Figure S4. IRF3 mRNA change was insignificant following RUVBL1/2 KD.

Figure S5. RUVBL1 and RUVBL2 mRNAs were unchanged between expression of e1aWT and e1a C-terminal mutants.

## Acknowledgements

This research was supported by the QCBio Collaboratory Fellowship 2019/2020 from the Institute for Quantitative & Computational Biosciences at the University of California, Los Angeles (K.E.K-U), and by the Professor June Lascelles Fund to the Department of Microbiology, Immunology and Molecular Genetics, UCLA.

## SUPPLEMENTARY METHODS AND MATERIALS

### siRNA and Small Molecule Treatment

siRNA KD was performed in HBTECs using Invitrogen RNAiMAX reverse transfection protocol. Cells were plated in antibiotic free HBTECs media containing indicated Ambion/ThermoFisher Silencer Select siRNA for a final concentration of 10nM that was pre-incubated in 7.5uL of lipofectamine RNAiMAX (ThermoFisher Cat#13778075) reagent in 750uL of Opti-MEM in 6cm^2^ plates. For a complete list of Ambion/ThermoFisher Silencer Select siRNAs used see below. Ambion/ThermoFisher Silencer Negative Control no.1 AM4611 was used as a negative control (siCtrl). Double knock-downs were performed with a total siRNA concentration of 10nM. MLN4924 (BostonBiochem Cat# I-502) was added directly to media of cultured HBTECs at a final concentration of 20µM for a treatment time of 6h. A485 (MedChemExpress Cat# HY-107455) was added directly to media of cultured HBTECs at a final concentration of 10µM for a treatment time of 12h. DMSO was used as a negative control for both MLN4924 and A485 at an equal volume.

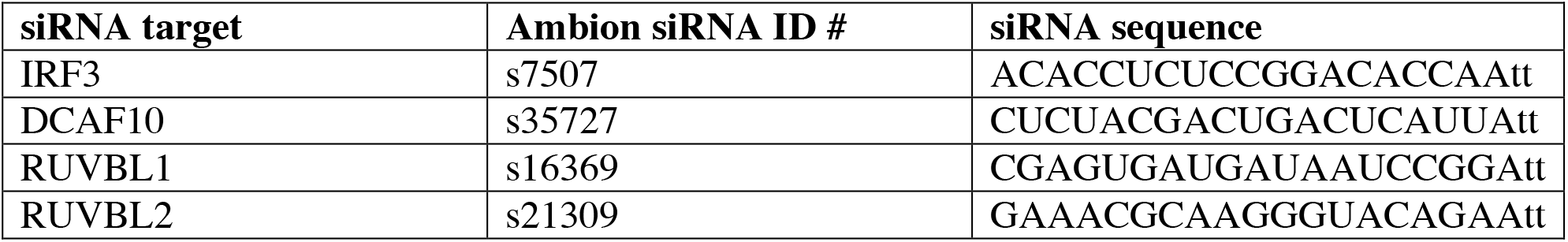

### Western Blot

Proteins were extracted from indicated cells by lysis in EBC (120 mM NaCl, 0.5% NP-40, 50 mM Tris-Cl pH 8.0, and Roche cOmplete protease inhibitors Cat#04693132001). Protein concentration was quantified by Bradford assay and normalized in Laemmli buffer and heated for 10min at 65°C then resolved in a 9% SDS-polyacrylamide gel. Proteins were electrotransferred to a polyvinylidene difluoride (PVDF) membrane then blocked in 5% milk in TBS-Tween 0.1% (blocking buffer) for 30 minutes. Primary antibody M58 (anti-E1A), M73 (anti-E1A), H3K27ac, H3, KU-86 (H-300), β-actin, IRF3, RUVBL1, RUVBL2, HUWE1, or DNAJA1 was added at 1:2000 dilution or 1:200 for M58 and M73 hybridoma supernatant for 1h at room temperature or O/N at 4°C. Membranes were washed 3X in TBS-Tween (0.1%) then HRP conjugated anti-mouse or anti-rabbit secondary antibodies were added for 1h room temperature in blocking buffer. Membranes were then washed 3X in TBS-Tween (0.1%) prior to addition of ECL reagent for detection of chemiluminescence. Western blots were validated with replicates of two or more with representative western blots presented.

| **Antibody** | **Source** | **Identifier** |
| --- | --- | --- |
| Rabbit polyclonal anti-H3K27ac | Active Motif | Cat#39133; RRID: AB_2561016; Lot#31814008 |
| Mouse monoclonal anti-E1A (M58) hybridoma supernatant | Produced in house | (30) |
| Rabbit polyclonal anti-Beta-Actin | GeneTex | Cat#GTX16039; RRID: AB_367276 |
| Rabbit polyclonal anti-KU-86 (H-300) | Santa Cruz | Cat#sc-9034; RRID: AB_2218743 |
| Rabbit monoclonal anti-IRF3 (D6I4C) | Cell Signaling Technology | Cat#11904S |
| Mouse monoclonal anti-H3 | Abcam | Cat#ab10799 |
| Anti-HA Tag (C29F4) | Cell Signaling Technology | Cat#3724S |
| Rabbit monoclonal anti-Pontin/RUVBL1 (D1L8J) | Cell Signaling Technology | Cat#74775S |
| Rabbit monoclonal anti-Reptin/RUVBL2 (D8N3J) | Cell Signaling Technology | Cat#12668S |
| Rabbit polyclonal anti-HUWE1 | Bethyl | Cat#A300-486A |
| Rabbit polyclonal anti-DNAJA1 | Bethyl | Cat#A304-516A |

### qRT-PCR

Cells were collected at indicated times following transfection and RNA was isolated using QIAGEN RNeasy Plus Mini Kit (Cat#74134). 1ug of RNA, as measured by Qubit fluorometer, was used for reverse transcription with SuperScript III First-Strand Synthesis SuperMix using random hexamer primers. qRT-PCR was performed with 5uL of cDNA, diluted 1:10. Runs were done using an ABI 7500 Real Time Thermocycler and reactions took place in optical-grade, 96-well plates (Applied Biosystems, Carlsbad, CA, USA) 25uL total volume with primers at a concentration of 900nM and 12.5uL of 2X FastStart Universal SYBR Green Master (Rox) (Roche Cat#04913850001). Relative mRNA levels were calculated as 2^ΔCt^. Equal cDNA loading was confirmed with primers to 18s rRNA cDNA. Data are presented as average of 3 or more biological replicates ± standard deviation.

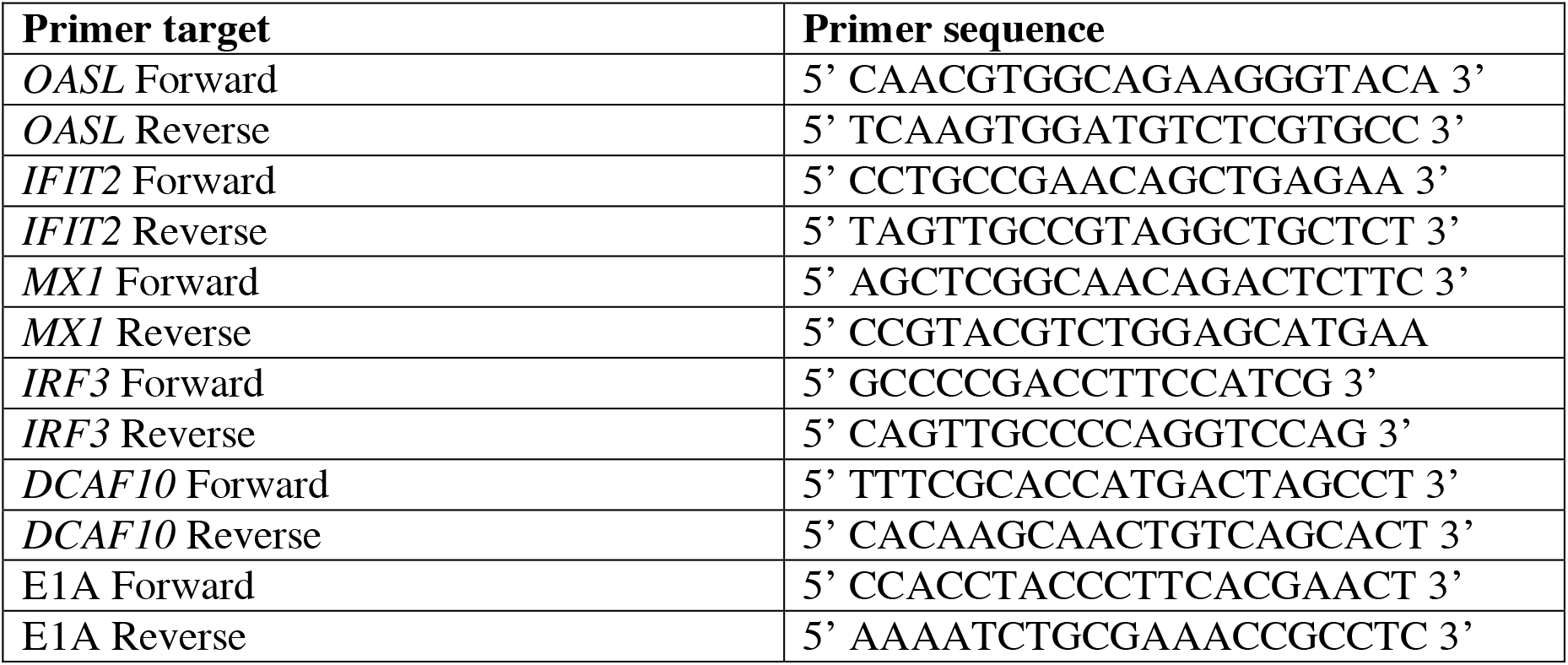

### RNA-seq Procedure and Data Analysis

1×10^6^ Low-passage HBTEC were mock-infected or infected with Ad5 E1A-E1B-substituted, E3-deleted vectors expressing WT Ad2 small E1A proteins from the dl1520 deletion removing the 13S E1A mRNA 50 splice site (*2*), 3 days after reaching confluence. RNA was isolated 24h p.i. using QIAGEN RNeasy Plus Mini Kit. Eluted RNA was treated with Ambion DNA-free™ DNA Removal Kit and then Ambion TRIzol reagent, precipitated with isopropanol, and dissolved in sterile water. RNA concentration was measured with a Qubit fluorometer. One microgram of RNA was fragmented and copied into DNA then PCR amplified with bar-coded primers for separate samples to prepare sequencing libraries using the Illumina TruSeq RNA Sample Preparation procedure. Libraries were sequenced using the Illumina HIseq-2000 to obtain single end 50-base-long reads. Sequences were aligned to the hg19 human genome sequence using TopHat v2. FPKM (fragments per kb per million mapped reads) for each annotated hg19 RefSeq gene ID was determined using Cuffdiff v2 from Cufflinks RNA-Seq analysis tools at http://cufflinks.cbcb.umd.edu. Homer (http://homer.salk.edu/homer PMID: 20513432) gene ontology enrichment analysis was performed on indicated gene lists. Homer motif discovery algorithm was used to look for transcription factor motifs +/-1 kb from the TSS of genes expressed 2X or more by all three e1a C-terminal mutants compared to e1aWT.

### Co-Immunoprecipitation

Co-IPs were performed using M58 crosslinked to protein G agarose beads or with goat anti-HA agarose immobilized (Bethyl cat#S190-138). 1mL of clarified M58 hybridoma supernatant was incubated with 50µL of 50% slurry protein G agarose beads on nutator for 4h at 4°C. Beads were washed 3X with 0.2M sodium borate pH 9 then antibody was crosslinked to protein G beads in 20mM DMP in 0.2M sodium borate pH 9 for 40 min on nutator at room temperature. Beads were then washed once with 0.2M ethanolamine pH 8 then quenched in 1mL ethanolamine pH 8 on nutator for 2h at room temperature. To remove uncoupled IgGs beads were washed 3X with 0.58% acetic acid and 150mM NaCl, then washed 3X with PBS.

Cells were lysed in EBC lysis buffer (120 mM NaCl, 0.5% NP-40, 50 mM Tris-Cl pH 8.0, and Roche cOmplete protease inhibitors and phosphatase inhibitors Sigma Aldrich cat# P5726-1ML and P0044-1ML) on ice. For western blots 2-4 mg of protein in supernatant lysate was precleared with 30µL agarose G beads for 1h then immunoprecipitated overnight at 4°C with M58 cross-linked to agarose G beads. Immuno-bead complexes were washed 3 times with cold EBC buffer and eluted in Laemmli buffer and incubated 10 min at 65°C. For mass spec samples 5-6 mg of protein in supernatant lysate was precleared with 50uL agarose G beads for 1h then 50uL of M58-beads or anti-HA beads were added for 4h at 4°C for immunoprecipitation. Immuno-bead complexes were washed 4 times with cold PBS then eluted in 500uL of 0.2M glycine pH 2.0. Proteins in 400uL of eluent were precipitated with 20% TCA on ice for 1h then microcentrifuged at 14K for 25 minutes at 4°C. Supernatant was removed and pellets were washed with 500uL of ice-cold acetone then microcentrifuged at 14K for 25 minutes at 4°C. Supernatant was removed and pellets were stored in parafilmed tubes at −20°C for eventual trypsin digest and mass spec.

### Sample Digestion and Desalting

The protein pellets were resuspended in digestion buffer (8M urea, 100mM Tris pH 8.5), later reduced and alkylated via sequential 20-minute incubations of 5 mM TCEP 10 mM iodoacetamide at room temperature in the dark while being mixed at 1200 rpm in an Eppendorf thermomixer. Then the proteins were digested by 0.1 μg of Lys-C (Thermo Scientific, 90051) and 0.8 μg Trypsin (Thermo Scientific, 90057) proteases at 37°C overnight. The digested samples were quenched by the addition of formic acid to 5% (v./v.). final concentration. Each sample was desalted via C18 stage tips (Thermo Scientific, 87784), washed twice in 200uL of 5% formic acid, and eluted in 40% Acetonitrile with 5% formic acid. Eluted peptide samples were dried in a SpeedVac before being resuspended in 5% formic acid.

### LC-MS Acquisition

Peptide samples were separated on a 75μM ID, 25cm C18 column packed with 1.9 μM C18 particles (Dr. Maisch GmbH HPLC) using a 140-minute gradient of increasing acetonitrile and eluted directly into a Thermo Orbitrap Fusion Lumos instrument where MS/MS spectra were acquired by Data Dependent Acquisition (DDA).

### Mass Spectrometry Analysis

Raw data for the e1a IP in A549 cells and HBTECs were converted into mzML format through the Proteowizard Package msconvert (1). Peptide spectrum matches were generated to identify fragmented peptide analytes through the MSGF+ search algorithm with a precursor ion tolerance of 20 ppm, allowing for isotopic error correction from −1 to +2 with the ‘addFeatures’ option enabled (2). Candidate peptides were considered from a fully tryptic digest of the EMBL human reference proteome, appended with the Adenovirus E1A protein sequence, between 6 to 40 amino acids in length. Peptide charge states were considered from +2 to +6, and a fixed modification of Carbamidomethylation was considered for Cysteine residues. Search outputs were converted to tabular results via msgf2pin, and the Percolator implementation included within the Crux package was used to assign confidence estimates to the peptide-spectrum matches and protein identifications (3–5). Peptide-spectrum matches and protein identifications were filtered at a maximum q-value of 0.01, and spectral counts for each acquisition tabulated via Crux’s ‘spectral-counts’ functionality.

For e1a IP experiments technical replicate data were produced by dividing material and performing two mass spec runs per condition. Data is presented as summed spectral counts for the duplicate runs or individual spectral counts for each technical replicate (Figure 2A).

Mass Spec analyses for HA-DCAF10 IP in A549 and HBTECs were performed by using ProLuCID (6) and DTASelect2 (7, 8) implemented in Integrated Proteomics Pipeline IP2 (Integrated Proteomics Applications). Protein and peptide identifications were filtered using DTASelect2 and required a minimum of two unique peptides per protein and a PSM-level false positive rate of less than 1% as estimated by a target-decoy database competition strategy. Protein identifications were filtered to require two uniquely mapping peptides to be considered confident.

## SUPPLEMENTARY FIGURES

**Figure S1.**
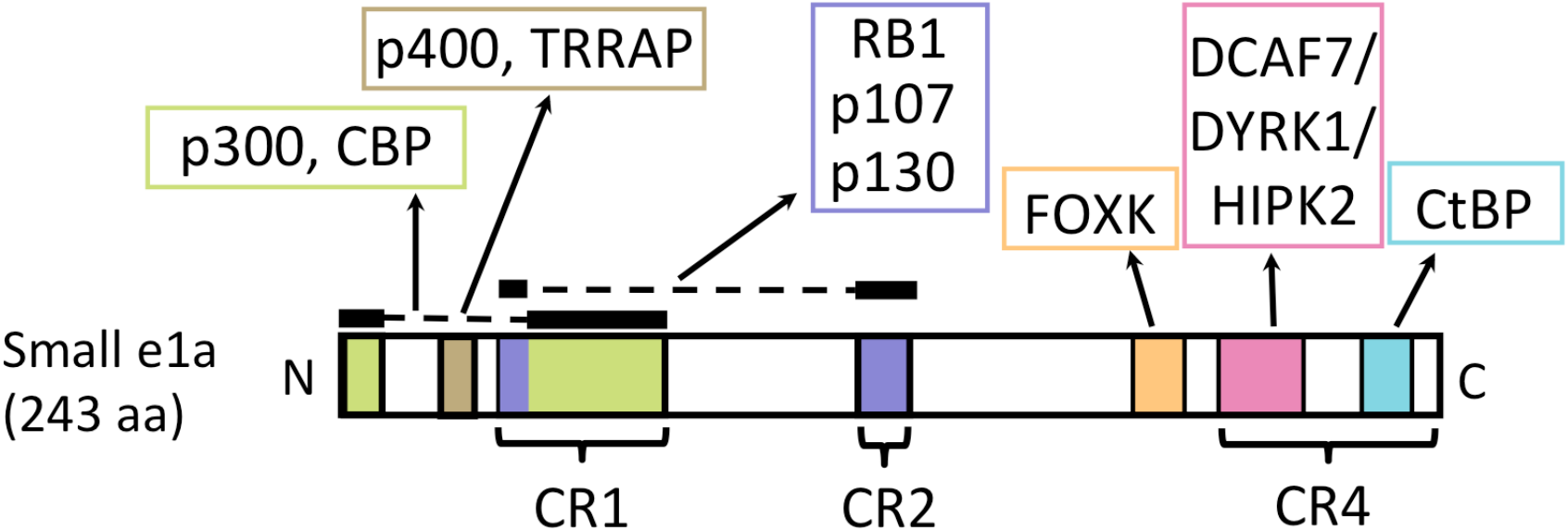
E1A regions bound by host proteins in primate adenoviruses. The N-terminal ∼15 aa residues plus the portion of CR1 diagrammed in green (aa 54-82) are intrinsically disordered protein (IDP) regions that fold and are bound by the p300/CBP TAZ2 domain (*14*), inhibiting p300/CBP histone acetyl transferase activity (*10, 15*). The N-terminal portion of CR1 (aa 37-49) and CR2 (aa 121-129, both blue) are bound by the “pocket” domains of RB-family proteins (*14, 16*). Between the N-terminal domain and CR1, aa 26-35 (brown region) mediate an interaction with p400 and TRRAP (*17*). The colored regions diagrammed in the C-terminal half of the E1A proteins are bound by FOXK1 and 2 (orange), the DCAF7 protein kinase complex (pink), and CtBP1 and 2 (light blue).

**Figure S2.**
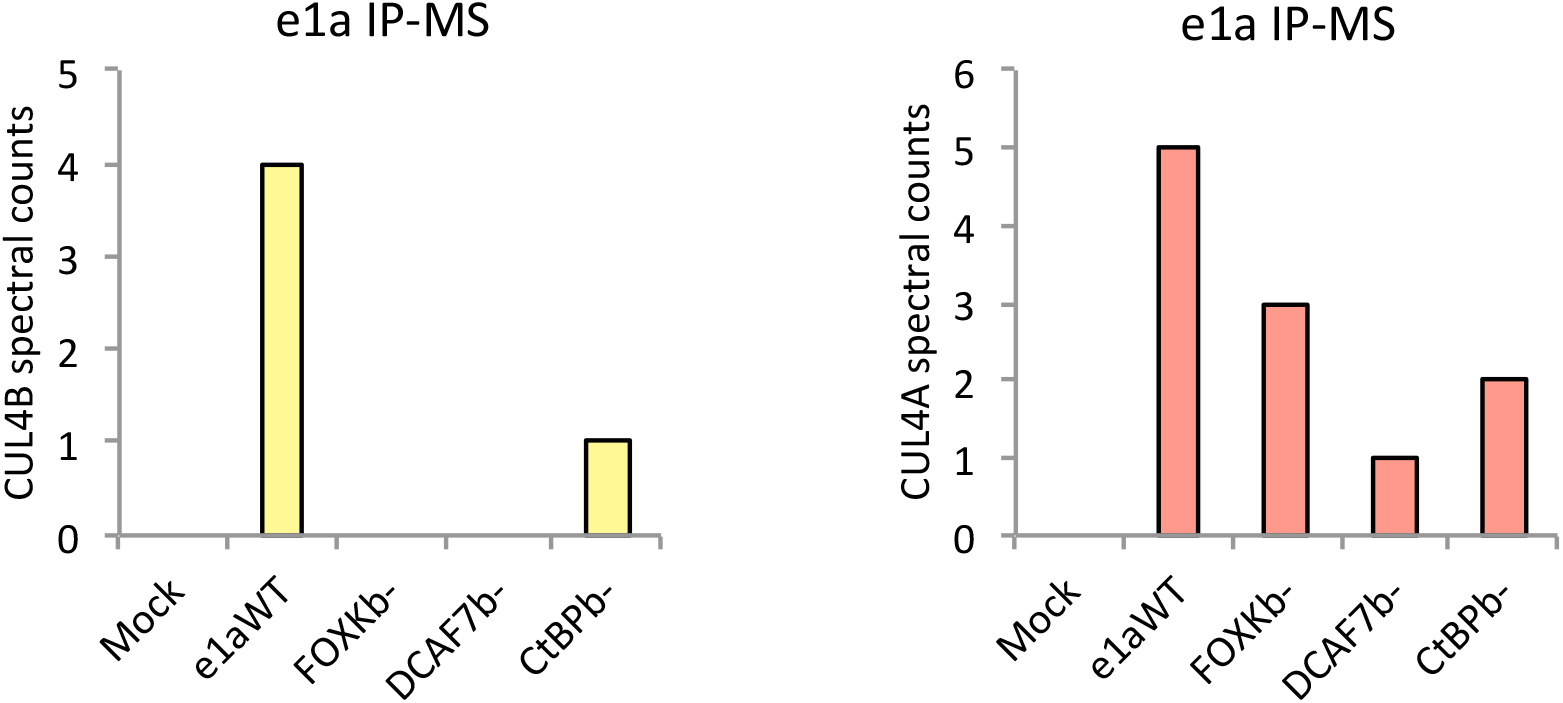
e1a C-terminal mutants have reduced binding to CUL4A/B. Spectral counts for CUL4B (left) and CUL4A (right) peptides in IPs with anti-e1a mAb M58 (anti-e1a) from extracts prepared 24h p.i. of A549 cells.

**Figure S3.**
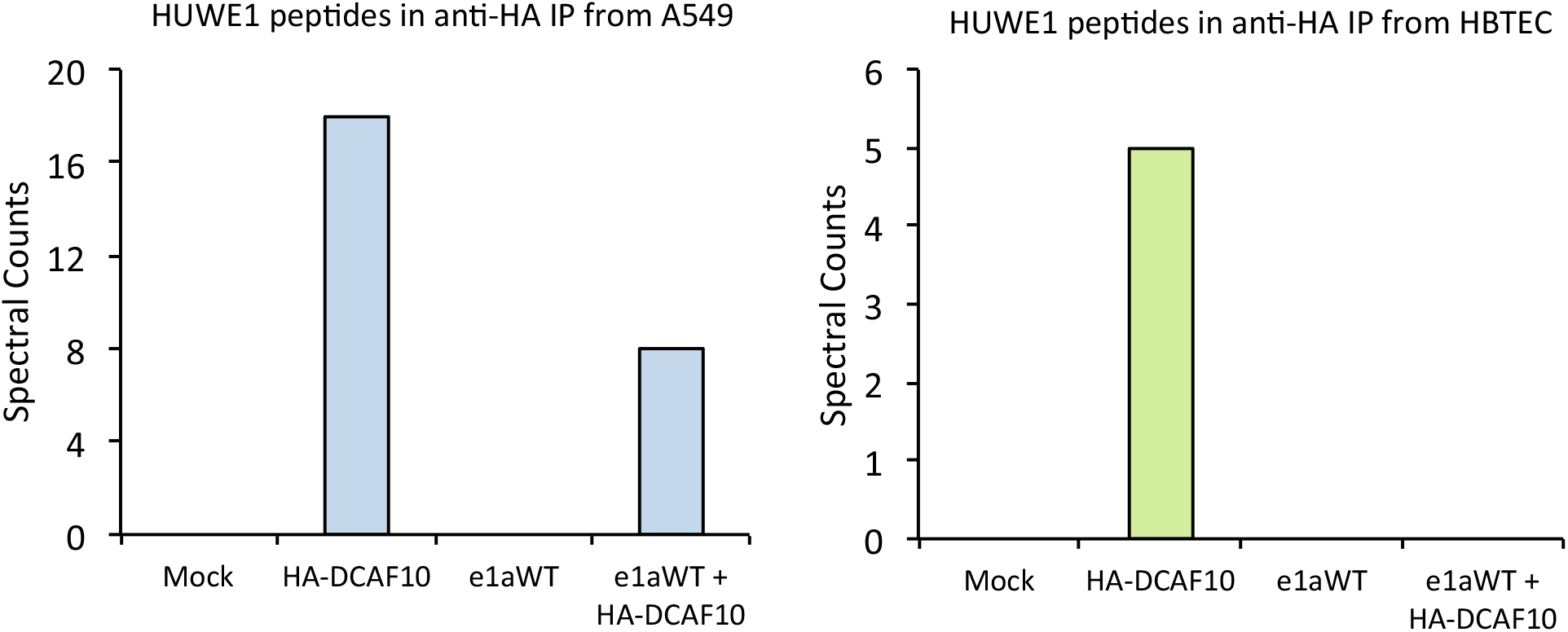
DCAF10 binds HUWE1. Spectral counts for HUWE1 mapped peptides from HA-DCAF10 (anti-HA) IP mass spec from 24h-infected A549 cells (left) and HBTECs (right).

**Figure S4.**
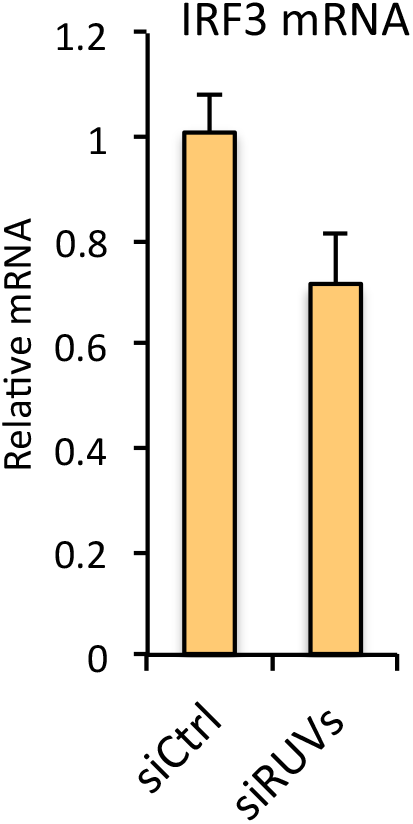
Insignificant change in IRF3 mRNA following RUVBL1/2 KD. Relative *IRF3* mRNA by qRT-PCR with cDNA synthesized from RNA extracted from three-day siRNA transfected HBTECs. siRUVs = treatment with siRNA against RUVBL1 and treatment with siRNA against RUVBL2. Data represent averages of three experimental replicates + SD.

**Figure S5.**
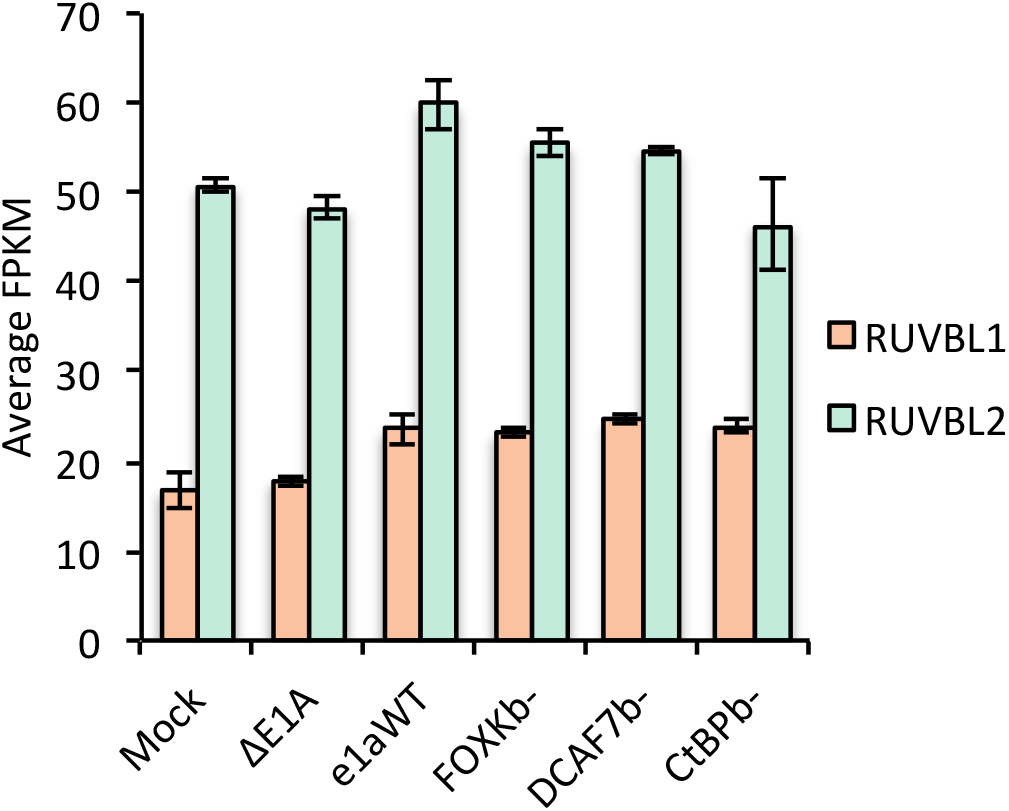
*RUVBL1* and *RUVBL2* mRNAs were unchanged between expression of e1aWT and e1a C-terminal mutants. FPKM of *RUVBL1* and *RUVBL2* from RNA-seq of 24h-infected HBTECs. Data represent averages of three experimental replicates + SD.

## References

1. G. Winberg, T. Shenk, Dissection of overlapping functions within the adenovirus type 5 E1A gene. EMBO J. 3, 1907–12 (1984).

2. C. Montell, G. Courtois, C. Eng, A. Berk, Complete transformation by adenovirus 2 requires both E1A proteins. Cell. 36, 951–961 (1984).

3. J. W. Lillie, M. R. Green, Transcription activation by the adenovirus E1a protein. Nature. 338, 39–44 (1989).

4. S. Stabel, P. Argos, L. Philipson, The release of growth arrest by microinjection of adenovirus E1A DNA. EMBO J. 4, 2329–2336 (1985).

5. H. E. Ruley, Adenovirus early region 1A enables viral and cellular transforming genes to transform primary cells in culture. Nature. 304, 602–606 (1983).

6. P. E. Branton, S. T. Bayley, F. L. Graham, Transformation by human adenoviruses. BBA - Rev. Cancer. 780 (1984), pp. 67–94.

7. M. Debbas, E. White, Wild-type p53 mediates apoptosis by E1A, which is inhibited by E1B. Genes Dev. 7, 546–554 (1993).

8. F. A. Dick, S. M. Rubin, Molecular mechanisms underlying RB protein function. Nat. Rev. Mol. Cell Biol. 14 (2013), pp. 297–306.

9. P. Pelka, J. N. G. Ablack, G. J. Fonseca, A. F. Yousef, J. S. Mymryk, MINIREVIEW Intrinsic Structural Disorder in Adenovirus E1A: a Viral Molecular Hub Linking Multiple Diverse Processes. J. Virol. 82, 7252–7263 (2008).

10. R. Ferrari, D. Gou, G. Jawdekar, S. A. Johnson, M. Nava, T. Su, A. F. Yousef, N. R. Zemke, M. Pellegrini, S. K. Kurdistani, A. J. Berk, Adenovirus small E1A employs the lysine acetylases p300/CBP and tumor suppressor Rb to repress select host genes and promote productive virus infection. Cell Host Microbe. 16, 663–676 (2014).

11. J. Komorek, M. Kuppuswamy, T. Subramanian, S. Vijayalingam, E. Lomonosova, L.-J. Zhao, J. S. Mymryk, K. Schmitt, G. Chinnadurai, Adenovirus type 5 E1A and E6 proteins of low-risk cutaneous beta-human papillomaviruses suppress cell transformation through interaction with FOXK1/K2 transcription factors. J. Virol. 84, 2719–2731 (2010).

12. M. J. Cohen, A. F. Yousef, P. Massimi, G. J. Fonseca, B. Todorovic, P. Pelka, A. S. Turnell, L. Banks, J. S. Mymryk, Dissection of the C-terminal region of E1A redefines the roles of CtBP and other cellular targets in oncogenic transformation. J. Virol. 87, 10348– 10355 (2013).

13. U. Schaeper, J. M. Boyd, S. Verma, E. Uhlmann, T. Subramanian, G. Chinnadurai, Molecular cloning and characterization of a cellular phosphoprotein that interacts with a conserved C-terminal domain of adenovirus E1A involved in negative modulation of oncogenic transformation. Proc. Natl. Acad. Sci. U. S. A. 92, 10467–10471 (1995).

14. J. C. Ferreon, M. A. Martinez-Yamout, H. J. Dyson, P. E. Wright, Structural basis for subversion of cellular control mechanisms by the adenoviral E1A oncoprotein. Proc. Natl. Acad. Sci. U. S. A. 106, 13260–13265 (2009).

15. E. Borrelli, R. Hen, P. Chambon, Adenovirus-2 E1A products repress enhancer-induced stimulation of transcription. Nature. 312, 608–612 (1984).

16. X. Liu, R. Marmorstein, Structure of the retinoblastoma protein bound to adenovirus E1A reveals the molecular basis for viral oncoprotein inactivation of a tumor suppressor. Genes Dev. 21, 2711–2716 (2007).

17. M. Fuchs, J. Gerber, R. Drapkin, S. Sif, T. Ikura, V. Ogryzko, W. S. Lane, Y. Nakatani, D. M. Livingston, The p400 complex is an essential E1A transformation target. Cell. 106, 297–307 (2001).

18. N. R. Zemke, A. J. Berk, The Adenovirus E1A C Terminus Suppresses a Delayed Antiviral Response and Modulates RAS Signaling. Cell Host Microbe. 22, 789-800.e5 (2017).

19. H. Negishi, T. Taniguchi, H. Yanai, The interferon (IFN) class of cytokines and the IFN regulatory factor (IRF) transcription factor family. Cold Spring Harb. Perspect. Biol. (2018), doi:10.1101/cshperspect.a028423.

20. H. Ikushima, H. Negishi, T. Taniguchi, The IRF family transcription factors at the interface of innate and adaptive immune responses. Cold Spring Harb. Symp. Quant. Biol. 78, 105–116 (2013).

21. N. Jones, T. Shenk, Isolation of adenovirus type 5 host range deletion mutants defective for transformation of rat embryo cells. Cell. 17, 683–689 (1979).

22. U. F. Greber, J. W. Flatt, Adenovirus Entry: From Infection to Immunity. Annu. Rev. Virol. 6, 177–197 (2019).

23. P. Ostapchuk, M. Suomalainen, Y. Zheng, K. Boucke, U. F. Greber, P. Hearing, The adenovirus major core protein VII is dispensable for virion assembly but is essential for lytic infection. PLoS Pathog. (2017), doi:10.1371/journal.ppat.1006455.

24. J. Chen, N. Morral, D. A. Engel, Transcription releases protein VII from adenovirus chromatin. Virology (2007), doi:10.1016/j.virol.2007.08.012.

25. M. M. Hu, H. B. Shu, Innate Immune Response to Cytoplasmic DNA: Mechanisms and Diseases. Annu. Rev. Immunol. (2020), doi:10.1146/annurev-immunol-070119-115052.

26. L. Gnatovskiy, P. Mita, D. E. Levy, The Human RVB Complex Is Required for Efficient Transcription of Type I Interferon-Stimulated Genes. Mol. Cell. Biol. (2013), doi:10.1128/mcb.01562-12.

27. O. Olanubi, J. R. Frost, S. Radko, P. Pelka, Suppression of Type I Interferon Signaling by E1A via RuvBL1/Pontin. J. Virol. (2017), doi:10.1128/jvi.02484-16.

28. H. Cheon, G. R. Stark, Unphosphorylated STAT1 prolongs the expression of interferon-induced immune regulatory genes. Proc. Natl. Acad. Sci. U. S. A. 106, 9373–9378 (2009).

29. L. M. Lasko, C. G. Jakob, R. P. Edalji, W. Qiu, D. Montgomery, E. L. Digiammarino, T. M. Hansen, R. M. Risi, R. Frey, V. Manaves, B. Shaw, M. Algire, P. Hessler, L. T. Lam, T. Uziel, E. Faivre, D. Ferguson, F. G. Buchanan, R. L. Martin, M. Torrent, G. G. Chiang, K. Karukurichi, J. W. Langston, B. T. Weinert, C. Choudhary, P. De Vries, J. H. Van Drie, D. McElligott, E. Kesicki, R. Marmorstein, C. Sun, P. A. Cole, S. H. Rosenberg, M. R. Michaelides, A. Lai, K. D. Bromberg, Discovery of a selective catalytic p300/CBP inhibitor that targets lineage-specific tumours. Nature. 550, 128–132 (2017).

30. E. Harlow, B. R. Franza, C. Schley, Monoclonal antibodies specific for adenovirus early region 1A proteins: extensive heterogeneity in early region 1A products. J. Virol. 55, 533–546 (1985).

31. S. Vijayalingam, T. Subramanian, L. Zhao, G. Chinnadurai, The Cellular Protein Complex Associated with a Transforming Region of E1A Contains c-MYC. J. Virol. 90, 1070–1079 (2016).

32. R. J. A. Grand, A. S. Turnell, G. G. F. Mason, W. Wang, A. E. Milner, J. S. Mymryk, S. M. Rookes, A. J. Rivett, P. H. Gallimore, Adenovirus early region 1A protein binds to mammalian SUG1-a regulatory component of the proteasome. Oncogene. 18, 449–458 (1999).

33. K. Burleigh, J. H. Maltbaek, S. Cambier, R. Green, M. Gale, R. C. James, D. B. Stetson, Human DNA-PK activates a STING-independent DNA sensing pathway. Sci. Immunol. 5 (5), doi:10.1126/sciimmunol.aba4219.

34. F. Glenewinkel, M. J. Cohen, C. R. King, S. Kaspar, S. Bamberg-Lemper, J. S. Mymryk, W. Becker, The adaptor protein DCAF7 mediates the interaction of the adenovirus E1A oncoprotein with the protein kinases DYRK1A and HIPK2. Sci. Rep. 6, 28241 (2016).

35. J. Jin, E. E. Arias, J. Chen, J. W. Harper, J. C. Walter, A Family of Diverse Cul4-Ddb1-Interacting Proteins Includes Cdt2, which Is Required for S Phase Destruction of the Replication Factor Cdt1. Mol. Cell. 23, 709–721 (2006).

36. T. A. Soucy, P. G. Smith, M. A. Milhollen, A. J. Berger, J. M. Gavin, S. Adhikari, J. E. Brownell, K. E. Burke, D. P. Cardin, S. Critchley, C. A. Cullis, A. Doucette, J. J. Garnsey, J. L. Gaulin, R. E. Gershman, A. R. Lublinsky, A. McDonald, H. Mizutani, U. Narayanan, E. J. Olhava, S. Peluso, M. Rezaei, M. D. Sintchak, T. Talreja, M. P. Thomas, T. Traore, S. Vyskocil, G. S. Weatherhead, J. Yu, J. Zhang, L. R. Dick, C. F. Claiborne, M. Rolfe, J. B. Bolen, S. P. Langston, An inhibitor of NEDD8-activating enzyme as a new approach to treat cancer. Nature. 458, 732–736 (2009).

37. S. P. Yamamoto, K. Okawa, T. Nakano, K. Sano, K. Ogawa, T. Masuda, Y. Morikawa, Y. Koyanagi, Y. Suzuki, Huwe1, a novel cellular interactor of Gag-Pol through integrase binding, negatively influences HIV-1 infectivity. Microbes Infect. 13, 339–349 (2011).

38. Y. Q. Mao, W. A. Houry, The role of pontin and reptin in cellular physiology and cancer Etiology. Front. Mol. Biosci. 4 (4), doi:10.3389/fmolb.2017.00058.

39. R. Lin, C. Heylbroeck, P. M. Pitha, J. Hiscott, Virus-Dependent Phosphorylation of the IRF-3 Transcription Factor Regulates Nuclear Translocation, Transactivation Potential, and Proteasome-Mediated Degradation. Mol. Cell. Biol. 18, 2986–2996 (1998).

40. H. Zimmermann, R. Degenkolbe, H.-U. Bernard, M. J. O’Connor, The Human Papillomavirus Type 16 E6 Oncoprotein Can Down-Regulate p53 Activity by Targeting the Transcriptional Coactivator CBP/p300. J. Virol. 73, 6209–6219 (1999).

41. R. Eckner, J. W. Ludlow, N. L. Lill, E. Oldread, Z. Arany, N. Modjtahedi, J. A. DeCaprio, D. M. Livingston, J. A. Morgan, Association of p300 and CBP with simian virus 40 large T antigen. Mol. Cell. Biol. 16, 3454–3464 (1996).

42. M. Nemethova, E. Wintersberger, Polyomavirus Large T Antigen Binds the Transcriptional Coactivator Protein p300. J. Virol. 73, 1734–1739 (1999).

43. E. Moran, M. B. Mathews, Multiple functional domains in the adenovirus E1A gene. Cell. 48 (1987), pp. 177–178.

44. H. G. Wang, E. Moran, P. Yaciuk, E1A promotes association between p300 and pRB in multimeric complexes required for normal biological activity. J. Virol. 69, 7917–24 (1995).

45. R. W. Stein, M. Corrigan, P. Yaciuk, J. Whelan, E. Moran, Analysis of E1A-mediated growth regulation functions: binding of the 300-kilodalton cellular product correlates with E1A enhancer repression function and DNA synthesis-inducing activity. J. Virol. 64, 4421–4427 (1990).

46. A. S. Turnell, R. J. Grand, C. Gorbea, X. Zhang, W. Wang, J. S. Mymryk, P. H. Gallimore, Regulation of the 26S proteasome by adenovirus E1A. EMBO J. 19, 4759– 4773 (2000).

47. M. Bouhamdan, S. Benichou, F. Rey, J. M. Navarro, I. Agostini, B. Spire, J. Camonis, G. Slupphaug, R. Vigne, R. Benarous, J. Sire, Human immunodeficiency virus type 1 Vpr protein binds to the uracil DNA glycosylase DNA repair enzyme. J. Virol. 70, 697–704 (1996).

48. Lynham J., Houry W.A. (2018) The Multiple Functions of the PAQosome: An R2TP- and URI1 Prefoldin-Based Chaperone Complex. In: Djouder N. (eds) Prefoldins: the new chaperones. Advances in Experimental Medicine and Biology, vol 1106. Springer, Cham. https://doi.org/10.1007/978-3-030-00737-9_4

49. M. I. Dauden, A. López-Perrote, O. Llorca, RUVBL1–RUVBL2 AAA-ATPase: a versatile scaffold for multiple complexes and functions. Curr. Opin. Struct. Biol. (2020) 67:78–85. doi:10.1016/j.sbi.2020.08.010.

50. E. Rajendra, J. I. Garaycoechea, K. J. Patel, L. A. Passmore, Abundance of the Fanconi anaemia core complex is regulated by the RuvBL1 and RuvBL2 AAA+ ATPases. Nucleic Acids Res. 42, 13736–13748 (2014).

51. M. A. Wood, S. B. McMahon, M. D. Cole, An ATPase/helicase complex is an essential cofactor for oncogenic transformation by c-Myc. Mol. Cell. 5, 321–330 (2000).

52. B. M. Iritani, R. N. Eisenman, c-Myc enhances protein synthesis and cell size during B lymphocyte development. Proc. Natl. Acad. Sci. U. S. A. 96, 13180–13185 (1999).

53. F. Martino, M. Pal, H. Muñoz-Hernández, C. F. Rodríguez, R. Núñez-Ramírez, D. Gil-Carton, G. Degliesposti, J. M. Skehel, S. M. Roe, C. Prodromou, L. H. Pearl, O. Llorca, RPAP3 provides a flexible scaffold for coupling HSP90 to the human R2TP co-chaperone complex. Nat. Commun. 9 (9), doi:10.1038/s41467-018-03942-1.

54. Z. Hořejší, H. Takai, C. A. Adelman, S. J. Collis, H. Flynn, S. Maslen, J. M. Skehel, T. de Lange, S. J. Boulton, CK2 phospho-dependent binding of R2TP complex to TEL2 is essential for mTOR and SMG1 stability. Mol. Cell. 39, 839–850 (2010).

55. S. Hardy, M. Kitamura, T. Harris-Stansil, Y. Dai, M. L. Phipps, Construction of adenovirus vectors through Cre-lox recombination. J. Virol. 71, 1842–1849 (1997).

## Supplemental References

1. D. Kessner, M. Chambers, R. Burke, D. Agus, P. Mallick, ProteoWizard: Open source software for rapid proteomics tools development. Bioinformatics (2008), doi:10.1093/bioinformatics/btn323.

2. S. Kim, N. Gupta, P. A. Pevzner, Spectral probabilities and generating functions of tandem mass spectra: A strike against decoy databases. J. Proteome Res. (2008), doi:10.1021/pr8001244.

3. L. Käll, J. D. Canterbury, J. Weston, W. S. Noble, M. J. MacCoss, Semi-supervised learning for peptide identification from shotgun proteomics datasets. Nat. Methods (2007), doi:10.1038/nmeth1113.

4. S. McIlwain, K. Tamura, A. Kertesz-Farkas, C. E. Grant, B. Diament, B. Frewen, J. J. Howbert, M. R. Hoopmann, L. Käll, J. K. Eng, M. J. MacCoss, W. S. Noble, Crux: Rapid open source protein tandem mass spectrometry analysis. J. Proteome Res. (2014), doi:10.1021/pr500741y.

5. V. Granholm, S. Kim, J. C. F. Navarro, E. Sjölund, R. D. Smith, L. Käll, Fast and accurate database searches with MS-GF +percolator. J. Proteome Res. (2014), doi:10.1021/pr400937n.

6. T. Xu, S. K. Park, J. D. Venable, J. A. Wohlschlegel, J. K. Diedrich, D. Cociorva, B. Lu, L. Liao, J. Hewel, X. Han, C. C. L. Wong, B. Fonslow, C. Delahunty, Y. Gao, H. Shah, J. R. Yates, ProLuCID: An improved SEQUEST-like algorithm with enhanced sensitivity and specificity. J. Proteomics (2015), doi:10.1016/j.jprot.2015.07.001.

7. D. L. Tabb, W. H. McDonald, J. R. Yates, DTASelect and contrast: Tools for assembling and comparing protein identifications from shotgun proteomics. J. Proteome Res. (2002), doi:10.1021/pr015504q.

8. D. Cociorva, D. L. Tabb, J. R. Yates, Validation of Tandem Mass Spectrometry Database Search Results Using DTASelect. Curr. Protoc. Bioinforma. (2007), doi:10.1002/0471250953.bi1304s16.

